# The development of the adult nervous system in the annelid *Owenia fusiformis*

**DOI:** 10.1101/2023.11.14.567050

**Authors:** Allan M. Carrillo-Baltodano, Rory Donnellan, Elizabeth A. Williams, Gáspar Jékely, José M. Martín-Durán

## Abstract

**Background:** The evolutionary origins of animal nervous systems remain contentious because we still have a limited understanding of neural development in most major animal clades. Annelids — a species-rich group with centralised nervous systems — have played central roles in hypotheses about the origins of animal nervous systems. However, most studies have focused on adults of deeply nested species in the annelid tree. Recently, *Owenia fusiformis* has emerged as an informative species to reconstruct ancestral traits in Annelida, given its phylogenetic position within the sister clade to all remaining annelids.

**Methods:** Combining immunohistochemistry of the conserved neuropeptides FVamide-lir, RYamide-lir, RGWamide-lir and MIP-lir with gene expression, we comprehensively characterise neural development from larva to adulthood in *Owenia fusiformis*.

**Results:** The early larval nervous system comprises a neuropeptide-rich apical organ connected through peripheral nerves to a prototroch ring and the chaetal sac. There are seven sensory neurons in the prototroch. A bilobed brain forms below the apical organ and connects to the ventral nerve cord of the developing juvenile. During metamorphosis, the brain compresses, becoming ring-shaped, and the trunk nervous system develops several longitudinal cords and segmented lateral nerves.

**Conclusions:** Our findings reveal the formation and reorganisation of the nervous system during the life cycle of *O. fusiformis*, an early-branching annelid. Despite its apparent neuroanatomical simplicity, this species has a diverse peptidergic nervous system, exhibiting morphological similarities with other annelids, particularly at the larval stages. Our work supports the importance of neuropeptides in animal nervous systems and the evolution of biphasic life cycles.

## Introduction

Nervous systems encompass all the neurons and their connections in an animal, representing an efficient way to communicate information along the body to elaborate behavioural and physiological responses in front of internal and external stimuli (1). Nervous systems are morphologically diverse, from diffuse nets as present in some non-bilaterian animals (e.g., ctenophores and cnidarians) to specialised and centralised systems with an anterior brain and post-cephalic longitudinal cords, as in many bilaterians (2, 3). Yet, how nervous systems evolved remains contentious because developmental information is lacking for many animal groups. Comparative, phylogenetically-guided studies on the specification, differentiation, patterning and architecture of nervous systems in as many different groups as possible (4, 5) are thus crucial to understand better how animal nervous systems originated and diversified (6).

Annelids — a group with a biphasic life cycle with a trochophore-like larva and centralised nervous systems as adults — have been central in understanding the evolution of nervous systems (3, 7–12). Traditionally, however, most studies have focused on species deeply nested in the annelid tree of life (13, 14), primarily on adults, and to a lesser extent using high-resolution developmental time courses (15–20). Therefore, studying lineages that branch off earlier in Annelida, such as Oweniidae, Magelonidae and Chaetopterimorpha, is essential to reconstruct ancestral traits in neural development for this animal clade (13, 21). Recent works in these groups (12, 21–26) have shown that a basiepidermal nervous system with a less organised brain was likely present in the last common annelid ancestor, which is a neuroanatomy that correlates well with their sedentary and tube-dwelling lifestyle (22). These studies have also indicated a simplification of the brain from larva to adult stages (22, 25, 26). However, we have previously demonstrated that the late embryos and early idiosyncratic mitraria larvae of the Oweniid *Owenia fusiformis* (23) show signs of organised neurogenesis in the anterior neural system where the apical organ forms and in the ciliary band that works as the main locomotory organ (24). With feeding, the mitraria larva undergoes a series of morphological transformations and increases in size (23, 24, 27, 28), concurrent to significant changes in gene regulation and the formation of a juvenile rudiment that broadly corresponds to the future adult trunk (23, 24, 27–29). However, using only a few immunostaining markers has prevented a better understanding of neural development in *O. fusiformis*, particularly during metamorphosis.

In this study, we combine cross-species antibodies against a variety of highly-conserved neuropeptides (30–32) with gene expression analyses of anterior marker genes (9, 33, 34) to characterise the development of the nervous system in *O. fusiformis*, from the larval to the adult stages (Figure 1). Our findings reveal a transition from a bilateral bilobed brain before metamorphosis that fuses during metamorphosis to give rise to a ring-shaped brain in the adult. Likewise, it provides new evidence of the brain’s connection with the future medullary cord of the trunk and the neural subdivisions in the segmented trunk. Together, we show a previously overlooked level of organisation of the nervous system in *O. fusiformis* that will be important to understanding the early dynamics of neural development in annelids and other animals.

**Figure 1.**
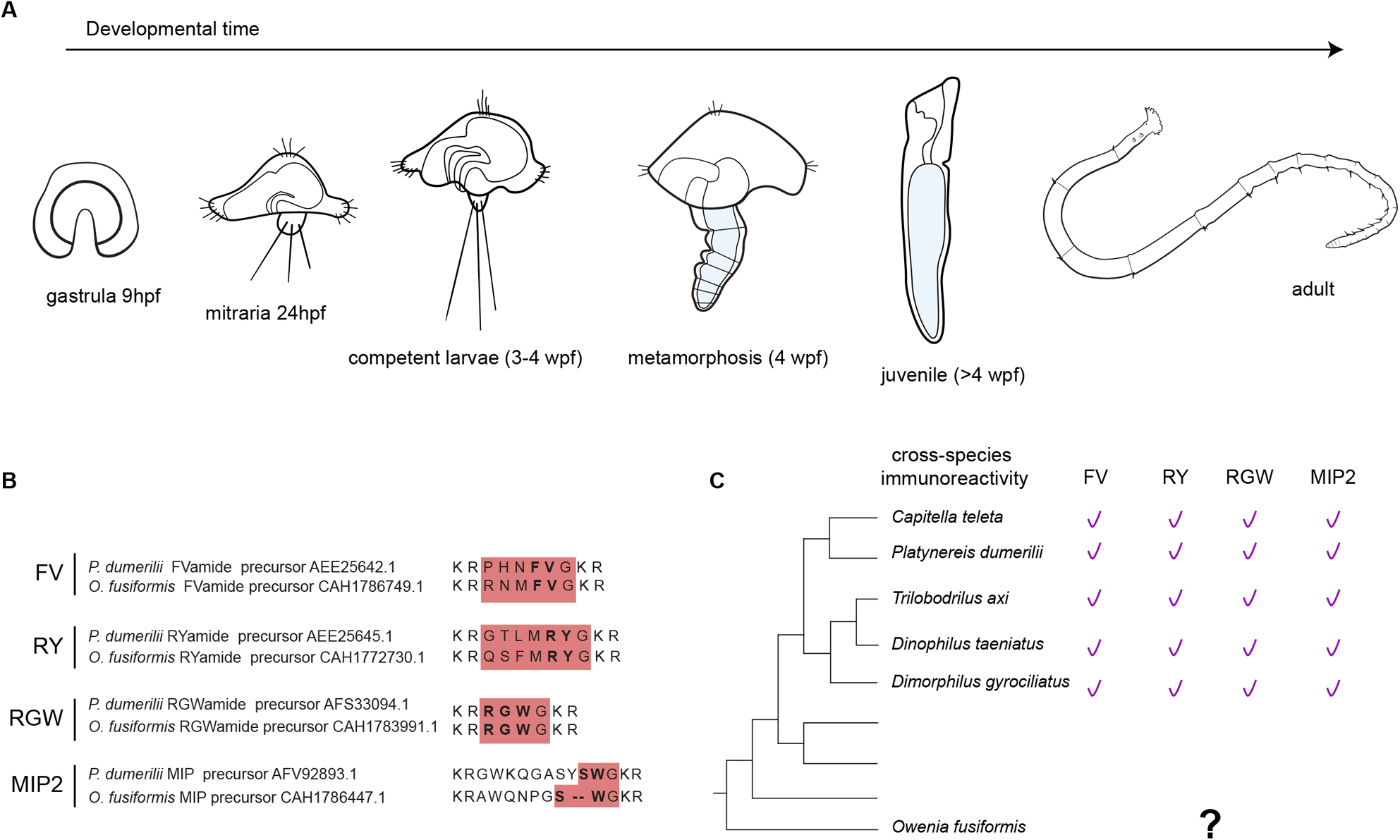
*Owenia fusiformis* development. **a** Developmental time course of stages studied: gastrula, early larva, competent larva, metamorphosis, juvenile and adult. **b** Conserved motifs in the epitopes of neuropeptides between *Platynereis dumerilii* (30–32) and *Owenia fusiformis*. Representative mature peptides and conserved dipeptides are highlighted in red and bold, respectively. **c** Cross-species reactivity tested across several annelids (30, 32, 40).

## Methods

### Animal collection

Reproductive individuals of *O. fusiformis* were collected from the coast near the Station Biologique de Roscoff (France) and kept in the laboratory as previously described (24, 33). Embryos and larvae were cultured as previously described (24).

### Immunohistochemistry

Fixation and antibody staining were conducted as described elsewhere (24). Adult specimens were relaxed in 8% MgCl_2_ and fixed overnight at 4°C. Adults were then placed in 60 mm dishes in 1x phosphate buffer saline (PBS), and their heads were dissected with a razor blade between the thoracic and the abdominal segments (between segments three and four (35, 36)). Adult heads were treated post-fixation with 1% collagenase D (Merk-Sigma, # COLLD-RO) overnight at 4°C and permeabilised through several washes with 1x PBS + 0.5% Triton X-100 (PTx). The primary antibodies mouse anti-acetylated α-tubulin (clone 6-11B-1, Merk-Sigma, #MABT868, 1:800), mouse beta-tubulin (E7, Developmental Studies Hybridoma Bank, 1:20), rabbit anti-FMRFamide (Immunostar, cat#: 20091, 1:600), and *Platynereis dumerilii* derived (30–32) rabbit anti-FVamide (stock concentration: 0.12 mg/ml; accession number: AEE25642.1, 1:200–1:500), anti-RYamide (stock concentration: 0.28 mg/ml; accession number: AEE25645.1, 1:200–1:500), anti-RGWamide (stock concentration: 0.4 mg/ml; accession number: AFS33094.1, 1:200–1:500) and anti-MIP (myoinhibitory peptide) (stock concentration: 0.28 mg/ml; accession number: AFV92893.1, 1:200–1:500) were diluted in 5% normal goat serum (NGS) in PTx and incubated overnight at 4°C. After several washes in 1% bovine serum albumin (BSA) in PTx, samples were incubated with secondary antibodies AlexaFluor488, AlexaFluor555 and AlexaFluor647 conjugated antibodies (ThermoFisher Scientific, A-21428, A32731, A-21235, 1:600) plus DAPI (stock 2mg/ml, 1:2000) diluted in 5% NGS in PTx overnight at 4°C. Adults were dehydrated stepwise in isopropanol, cleared in 2:1 benzyl benzoate:benzyl alcohol, briefly immersed in xylene, and mounted in Entellan (Merk-Sigma, #1.07960).

### Orthology analysis

A previously published alignment of SOX proteins (37) and maximum likelihood tree reconstruction with FastTree (38) were used to assign the orthology of SOXC in *O. fusiformis*.

### Whole-mount in situ hybridisation

Riboprobes were synthesised with the T7 enzyme following the manufacturer’s recommendations (Ambion’s MEGAscript kit, #AM1334) and stored in hybridisation buffer at a concentration of 50 ng/μl at −20°C. Single colourimetric *in situ* hybridisation of embryos and mitraria larvae was performed following an established protocol using a 1.5 ng/μl probe concentration (24, 29, 33, 34).

### Imaging

Representative embryos, larvae, and juveniles from the colourimetric whole mount *in situ* hybridisation experiments were cleared and mounted in 80% glycerol in PBS. They were imaged with a Leica DMRA2 upright microscope equipped with an Infinity5 camera (Lumenera) using differential interference contrast (DIC) optics. Confocal laser scanning microscopy (CLSM) images were taken with a Leica SP5, Leica Stellaris 8 and Nikon CSU-W1 spinning disk confocal microscope. CLSM Z-stack projections were built with ImageJ2 (39) and Nikon NIS-elements software. DIC images were digitally stacked with Helicon Focus 7 (HeliconSoft). Brightness and contrast were edited with Adobe Photoshop CC (v 24.0.0), and figures were built with Adobe Illustrator CC (v 27.0.0) (Adobe Inc.).

## Results

To characterise better the complexity and development of the nervous system of *O. fusiformis,* we tested four purified antibodies against conserved mature neuropeptides (FVamide, RYamide, RGWamide and MIP) of the annelid *P. dumerilii* that have broad cross-species immunoreactivity (Figure 1b–c; Figure 2; Additional File 1: Supplementary Figure 1; Additional File 2: Supplementary Figure 2) (30, 32, 40). FVamide, RYamide and RGWamide label many of the previously described components of the early larval nervous system (24) (Figure 2), including the apical organ and the prototroch ring, but also previously uncharacterised peripheral nerves in the larval episphere. The MIP antibody has a lower signal-to-noise ratio but still labels the apical organ and some tissue anterior to the larval mouth (Additional File 2: Supplementary Figure 2). Having confirmed their connection to the larval neural components, we focused on describing the immuno-reactivity of these antibodies during the life cycle of *O. fusiformis*, using tubulin as a counter-immunostaining of the nervous system.

**Figure 2.**
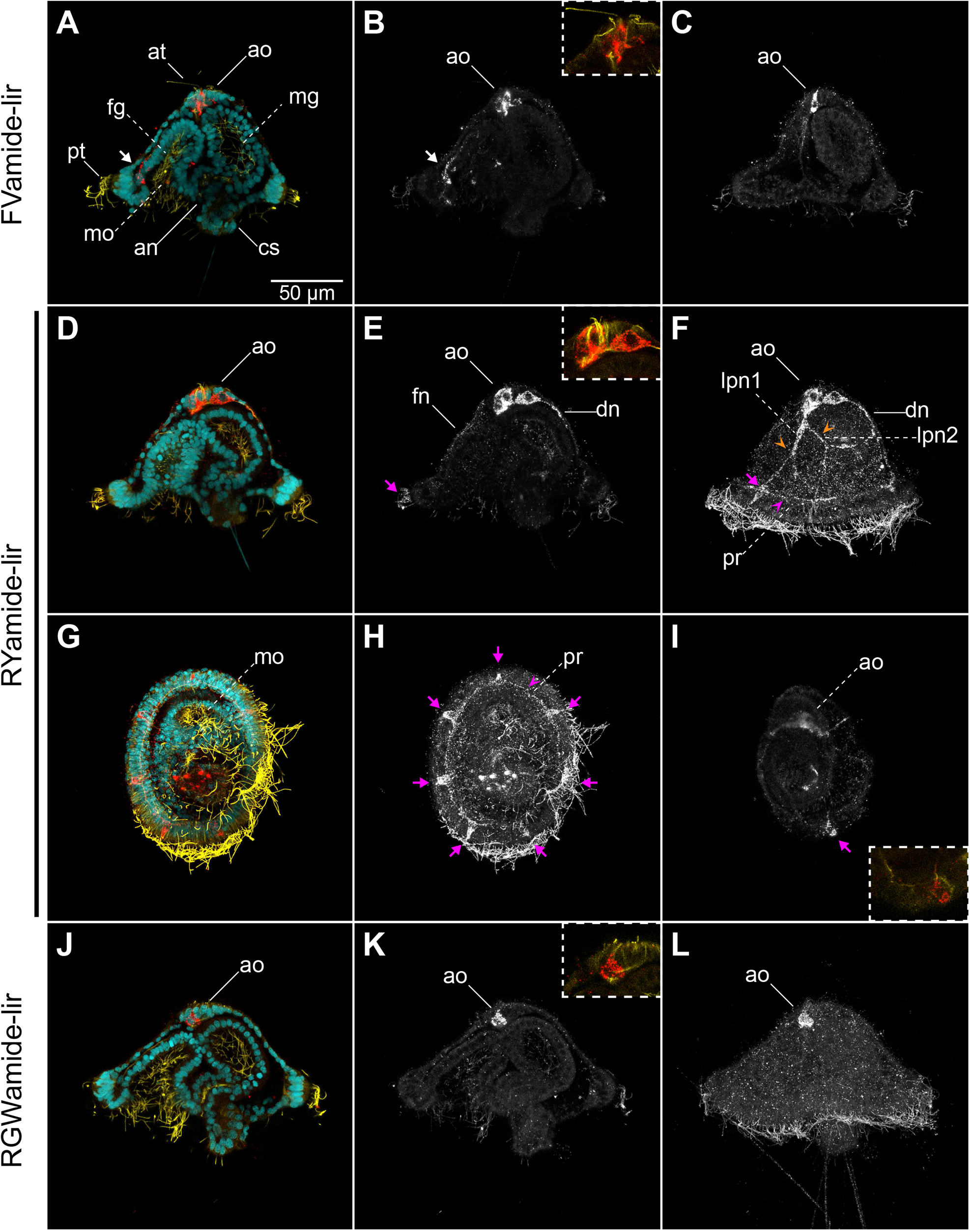
Neuropeptide-lir elements in the early mitraria. Confocal Laser Scanning Microscopy (CLSM) images of DAPI (cyan), acetylated tubulin (yellow) and neuropeptide-lir (red or white) elements at 24 hpf. All images are lateral views except for ventral views in **g**–**i**. Insets in (**b**, **e**, **i and k**) are close ups of the apical organ (ao) in the same view as the larger image. **a–c** FVamide-lir cells in the apical organ and one cell anterior to the foregut (white arrow). **d**–**i** RYamide-lir cells are present in the apical organ, with RYamide-lir axons (fn, dn, and orange arrowheads) connecting with seven RYamide-lir cells (magenta arrows) and an RYamide-lir prototrochal ring (pr). **j**–**l** RGWamide-lir cells are exclusively present in the apical organ. an: anus; ao: apical organ; at: apical tuft; cs: chaetal sac; dn: dorsal nerve; fg: foregut; fn: frontal nerve; mg: midgut; mo: mouth; pr: prototrochal ring; pt: prototroch.

### The complex nervous system of the early mitraria

At 24 hours post-fertilisation (hpf), between three to seven FVamide-like immune-reactive (FVamide-lir), RYamide-lir and RGWamide-lir cells are detectable in the apical organ of the early mitraria larva (Figure 2). A solitary FVamide-lir neuron with a weak FVamide-lir short axon is positioned anterior and apical to the mouth (white arrow, Figure 2a–b). MIP has a similar pattern of immunoreactivity (white arrow, Additional File 2: Supplementary Figure 2). RYamide-lir axons, on the other hand, connect the apical organ to an RYamide-lir prototroch ring (pr) (magenta arrowhead, Figure 2e–f, h) via a frontal nerve (fn), a dorsal nerve (dn) and two bilateral peripheral nerves (lpn1–2) that bifurcate further midway in the episphere (orange arrowheads, Figure 2e–f). The prototroch ring also contains seven RYamide-lir cells (magenta arrows, Figure 2e–f, h–i), three anterior and four posterior, similar to the FMRFamide-lir, *elav*^+^ and *synaptotagmin*^+^ cells previously described at this larval stage (24). In contrast, RGWamide-lir cells are exclusively restricted to the apical organ (Figure 2j–l). Apical cilia protrudes from some of the FVamide-lir, RYamide-lir, RGWamide-lir and MIP-lir neurons of the apical organ (Figure 2b, e, j; Additional File 2: Supplementary Figure 2). At this stage, beta-tubulin and alpha-acetylated tubulin label the frontal, dorsal, and peripheral nerves connecting the apical organ with the tubulin^+^ prototroch ring (Figure 3a–e, h–m). Near the seven refringent globules of unknown function (24, 27), but integrated within the prototroch, are at least five beta-tubulin^+^ monociliated cells with a short cilium, which likely represent mechanoreceptors (Figure 3e–g). Together, these new neuropeptide antibodies and more detailed observations of tubulin immunostaining demonstrate the complexity of the apical organ and neural components of the prototroch, including elaborated neurite patterns that connect these two sensorial structures, many of which had been previously overlooked (9, 23, 24).

**Figure 3.**
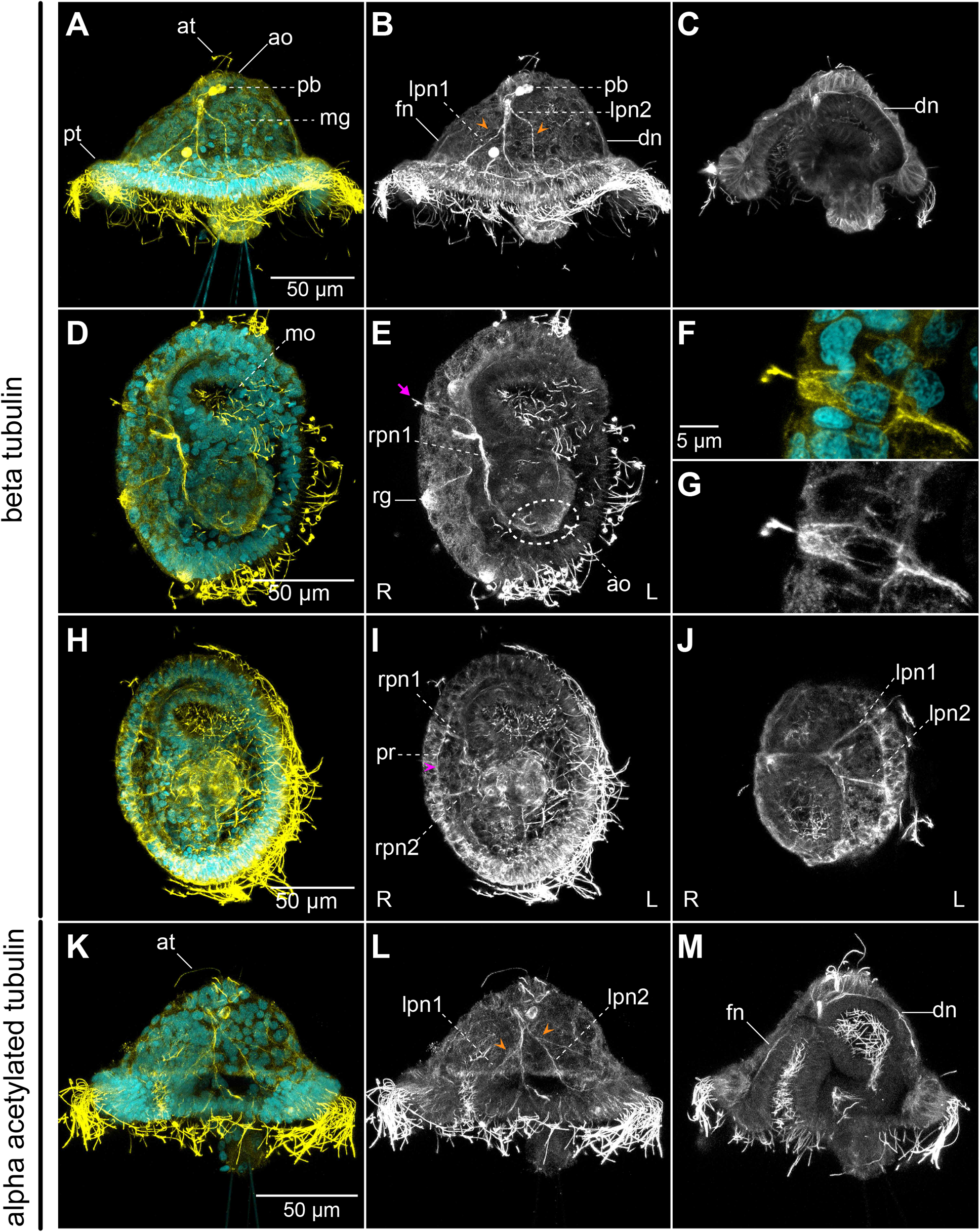
Tubulin^+^ elements in the early mitraria. CLSM images of DAPI (cyan) and beta-tubulin (**a**–**j**) and alpha-acetylated tubulin (**k**–**m**) (yellow) at 24 hpf. Insets in (**f**–**g**) are close ups of the peripheral neuron in **e**. **a**–**c** Lateral views with beta-tubulin^+^ axons extending from the apical organ (ao) anteriorly (fn), dorsally (dn) and laterally (rpn, lpn; orange arrowheads) towards the prototroch ring. The polar bodies (pb) are still visible at this stage in the blastocoel space between the apical organ and the midgut (mg). Beta-tubulin is also staining the cell boundaries across the body of the larva, like in **c**. **d**–**g** Ventral views showing at least one beta-tubulin^+^ monociliated cell (magenta arrow) in the prototroch that presumably connects to the apical organ via a peripheral nerve (rpn1). **h**–**j** Two bilateral peripheral nerves (rpn1–rpn2 and lpn1–lpn2) branch out on each side of the episphere towards the tubulin^+^ prototrochal ring (pr). **k**–**m** Most of the beta-tubulin^+^ axons are also with acetylated tubulin. ao: apical organ; at: apical tuft; dn: dorsal nerve; fn: frontal nerve; lpn1–lpn2: left peripheral nerves 1–2; mg: midgut; mo: mouth; pr: prototrochal ring; pt: prototroch; rg: refringent globule; rpn1–rpn2: right peripheral nerves 1–2.

### The formation of the brain and nerve cords

As the larva grows and acquires competence, the adult brain forms, first as a horseshoe-shaped, bilobular, apical condensation of nuclei recognisable, as well, through the cell membrane labelling with beta-tubulin (27) (br; Figure 4a, e, g, k, m, o, q, u; Additional File 3: Supplementary Figure 3a, d, g, j; Additional File 4: Supplementary Figure 4a–d). In addition, the bilateral gene expression of the putative neural gene *soxC* (Additional File 5: Supplementary Figure 5) and anterior markers *pou4*, *six3/6*, *nk2.1* and *ChAt* (Figure 5a–f, i–l) confirm the bilobular nature of the brain at this stage. A small pit, as referred to by Wilson (27), is positioned most apically in the brain, where the ciliated apical tuft protrudes (Additional File 4: Supplementary Figure 4a–b, g–i). In addition, an apical ring of FVamide-lir, RYamide-lir, MIP-lir, and tubulin^+^ cells surround this apical tuft (ar; Figure 4f, j, v; Additional File 4: Supplementary Figure 4b, h) and is presumably part of the apical organ. At this stage, this neural larval organ also contains multiple FVamide-lir, RYamide-lir, RGWamide-lir and MIP-lir neurons, interconnected with the brain sitting just below (ao; Figure 4 a–b, d–f, g–l, m–r, u–v; Additional File 3: Supplementary Figure 3). Two thick RYamide-lir and tubulin^+^ axon bundles — the ventral and dorsal roots — cross the brain and form a central neuropil just below the condensed nuclei of the brain (23, 27) (Figure 4k–l; Additional File 4: Supplementary Figure 4c, h). We used the terms “ventral” root and “dorsal” root to follow the nomenclature of the brain in other annelids (14, 41). However, the ventral and dorsal roots are positioned anteriorly and posteriorly, respectively, along the main body axis of the larva and juvenile. Altogether, these apical neural structures connect with the developing ventral nerve cord (vnc) of the juvenile rudiment (see below) through FVamide-lir, MIP-lir (Additional File 3: Supplementary Figure 3b, k), and tubulin^+^ (Additional File 4: Supplementary Figure 4b) circumesophageal connectives. Eyespots are present on each side of the most basal part of the brain (not shown) (23, 27). Lastly, frontal and dorsal nerves, plus the lateral peripheral nerves, maintain the connection between the apical organ/brain and the prototroch neural ring (Figure 4b, h, r; Additional File 3: Supplementary Figure 3b, e, k; Additional File 4: Supplementary Figure 4f, i).

**Figure 4.**
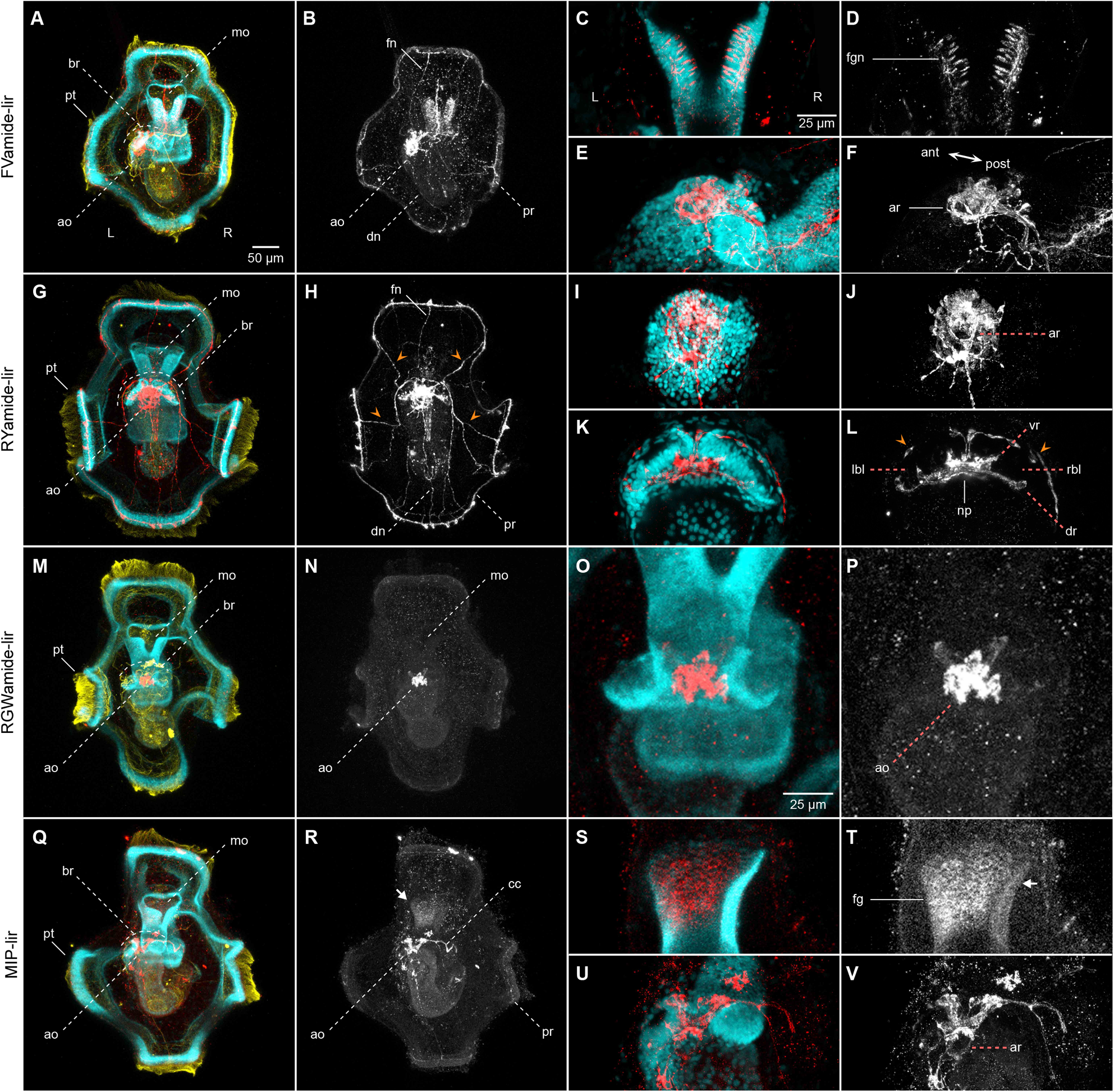
Neuropeptide-lir elements in the competent larvae. CLSM images of DAPI (cyan), acetylated tubulin (yellow) and neuropeptide-lir (red or white) elements in the competent larvae (∼ 3 wpf). Apical views, with anterior to the top. **c**–**f**, **i**–**l**, **o**–**p** and **s–v** are close ups of the foregut or apical organ in the same view as the respective larger image in **b**, **h**, **n**, **r**. **a**–**b**, **e**–**f** FVamide-lir cells and **q**–**r**, **u**–**v** MIP-lir cells in the apical organ connect via FVamide-lir and MIP circumesophageal connectives (cc) to the ventral nerve cord (vnc) of the juvenile trunk rudiment (jr) (See Additional File 3: Supplementary Figure 3), and via **a**–**b** FVamide-lir, **g**–**h** RYamide-lir and **q**–**r** MIP-lir frontal (fn), dorsal (dn) and peripheral nerves (orange arrow heads) to the **a**–**b** FVamide-lir, **g**–**h** RYamide-lir and **q**–**r** MIP-lir prototrochal ring (pr). An **e**–**f** FVamide-lir, **i**–**j** RYamide-lir and **u**–**v** MIP-lir apical nerve ring (ar) surrounds the apical tuft. The foregut is innervated by **a**–**d** FVamide-lir cells and neurites. **k**–**l** RYamide-lir axons form a neuropil between two brain lobes (rbl–lbl) underneath the apical organ. **m**–**p** RGWamide-lir cells remain only in the apical organ. Arrow in **r**, **t** is presumably background staining.an: anus; ao: apical organ; ar: apical nerve ring; at: apical tuft; br: brain; cc: circumesophageal connectives; chn: chaetal sac nerve; cs: chaetal sac; dn: dorsal nerve; dr: dorsal root; fg: foregut; fgn: foregut nerve; fn: frontal nerve; jr: juvenile rudiment; mg: midgut; mo: mouth; np: brain neuropil; pr: prototrochal ring; pt: prototroch; vr: ventral root.

**Figure 5.**
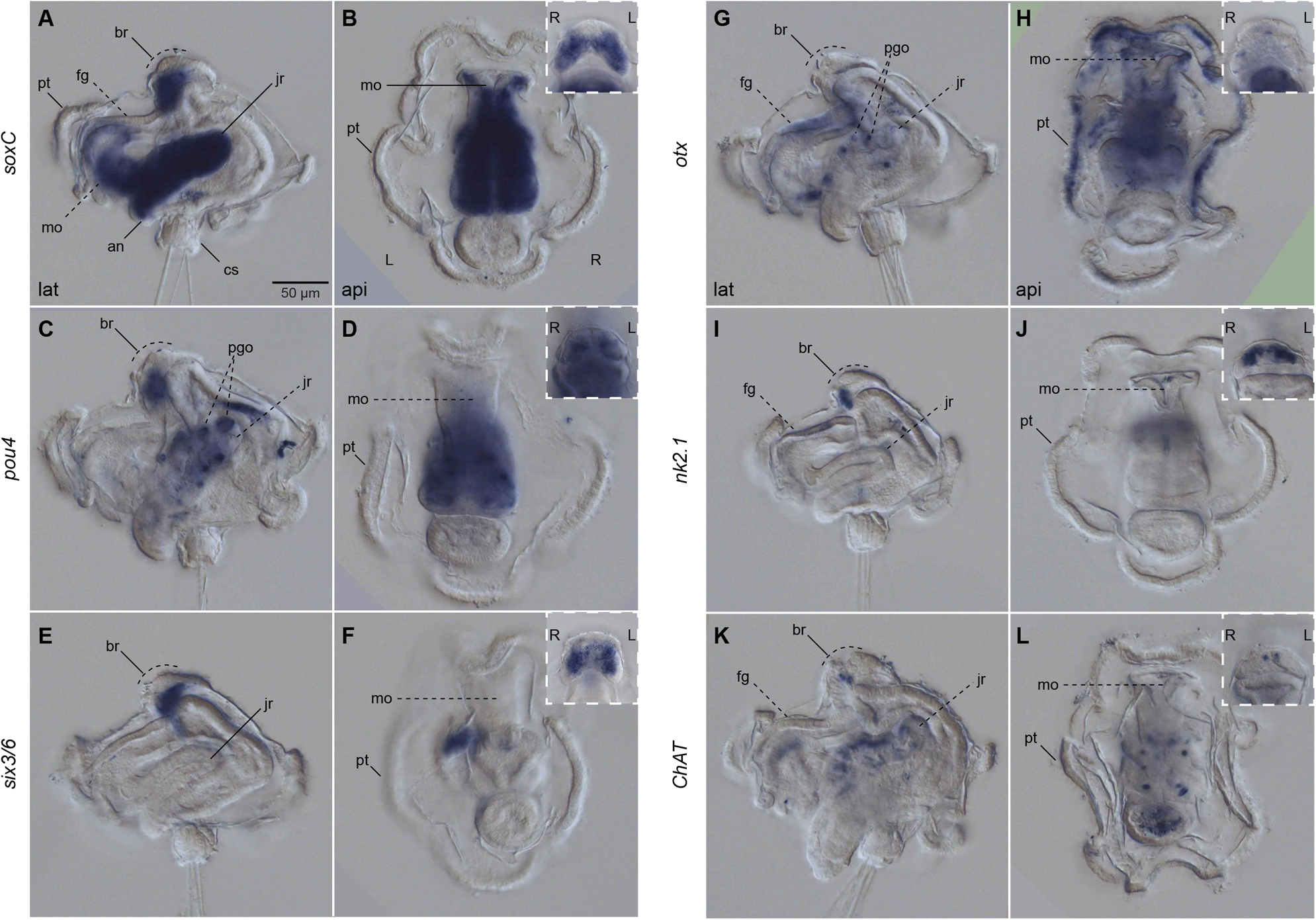
Expression of neural genes in the competent larvae. Differential Interference Contrast (DIC) images showing expression of *soxC*, *pou4*, *six3/6*, *otx*, *nk2.1* and *ChAt*. **a**, **c**, **e**, **g**, **i**, **k** lateral views; **b**, **d**, **f**, **h**, **j**, **l** apical views. Insets are close ups of the corresponding larger images in anterior view. All genes, except for *otx* **g**–**h**, have a bilateral expression in the brain (br). **a**–**b** *soxC* is strongly expressed in the juvenile rudiment (jr), the mouth (mo), and the anterior part of the foregut (fg). **g**–**l** *otx*, *nk2.1* and *ChAt* have some weaker expression in the foregut. **g**–**h** in addition *otx* is expressed in the prototroch. an: anus; br: brain; cs: chaetal sac; fg: foregut; jr: juvenile rudiment; mo: mouth; pgo: parapodial glandular organ; pt: prototroch.

At this pre-metamorphic larval stage, the juvenile rudiment has grown into a defined trunk, with segments that will wrap around the gut as it prepares to evaginate from the larval body (27–29). The vnc of the trunk starts forming as early as two weeks post fertilisation (wpf) and is immunoreactive to serotonin (5HT), FMRFamide and tubulin (23). Between two to three wpf, 5HT-lir and FMRFamide-lir neurons and lateral nerves presumably get patterned on each of the developing trunk segments (23). In agreement with the expression of *elav* and *synaptotagmin*, *soxC* is highly expressed in the juvenile trunk at this stage, supporting that this is a prominent site of active neurogenesis in the competent larva (24) (Figure 5a–b). Not only has the trunk an FVamide-lir, RYamide-lir, MIP-lir and tubulin^+^ vnc but also an FVamide-lir and RYamide-lir dorsal one (Additional File 3: Supplementary Figure 3c, f), demonstrating that many of the components of the adult peripheral nervous system develop before metamorphosis.

In addition to the developing brain and nerve cords, the foregut is innervated with FVamide-lir and RYamide-lir neurons and nerves (fgn; Figure 4d–d, h; Additional File 3: Supplementary Figure 3b–c, e–f). MIP shows some unspecific labelling at the anterior section of the foregut (Figure 4s–t; Additional File 3: Supplementary Figure 3k–l), mirroring the expression domains of *soxC*, *otx*, *nk2.1*, and *ChAt* in this larval region (Figure 5a, g, i, k). Dorsal to the posterior tip of the trunk, the larval chaetal sac, which has many more chaetae at this stage than in the early mitraria, has an RYamide-lir and MIP-lir nerve connecting these defensive structures to the peripheral neurites of the episphere (Additional File 3: Supplementary Figure 3f, l). Altogether, the comprehensive analysis of the nervous system of the competent larva of *O. fusiformis* reveals a transition of neural connectivity, where the forming adult brain remains connected to the transitory larval organs, such as the prototroch and chaetal sac, as the connections with the developing trunk nervous system are established.

### The nervous system during metamorphosis

The apical organ remains positioned dorsally and apically to the double root of axons of the brain (i.e., the central neuropil; Figure 6; Additional File 6: Supplementary Figure 6a), and continues to be connected with the larval episphere and prototroch ring with the FVamide-lir, RYamide-lir and tubulin^+^ dorsal nerves (Additional File 6: Supplementary Figure 6a–d; Additional File 7: Supplementary Figure 7b), and RYamide-lir and tubulin^+^ lateral nerves (Figure 6c–d; Additional File 6: Supplementary Figure 6c–d; Additional File 6: Supplementary Figure 7a–b). The distinct two lobes of the brain of the competent larva appear to fuse into a continuous horseshoe during metamorphosis (Figure 7b, f), forming the putative ring-shaped brain of the juvenile and adult (see below). The dorsal and ventral root of the brain creates an FVamide-lir, RYamide-lir, RGWamide-lir, MIP-lir and tubulin^+^ neuropil (np; Figure 6c, f, i, l; Additional File 7: Supplementary Figure 7a–b), which connects to the thorax of the evaginating trunk via circumesophageal connectives (or lateral medullary cords (22); see discussion) (Figure 6; Additional File 6: Supplementary Figure 6; Additional File 7: Supplementary Figure 7a–b). In the juvenile and adult, the thorax is composed of three fused trunk segments, which we name ciliated thoracic segments (cts), and differentiate from the other trunk segments by having capillary chaetae (35, 36) and abundant cilia in the epidermis (Additional File 7: Supplementary Figure 7a–b). Paired RGWamide-lir parapodial glandular organs (pgos) up to the seventh segment (27, 42) facilitate the distinction between the three thoracic and the seven abdominal segments (27, 28) (Figure 6g–h; Additional File 6: Supplementary Figure 6e–f). We could not observe ganglia in either thoracic or abdominal segments using nuclear staining and gene expression (Figure 7), providing further evidence of the medullary cord nature in oweniids (12, 22). However, several iterated FVamide-lir, RYamide-lir, RGWamide-lir, and MIP-lir neurons are present along the vnc, which are more condensed in the thorax because of the fusion of the three thoracic segments and more distant in the rest of the trunk (Figure 6a–b, d–e, g–h, j–k; Additional File 6: Supplementary Figure 6). From these clusters of iterated neurons, FVamide-lir, RYamide-lir, RGWamide-lir, MIP-lir and tubulin^+^ lateral nerves run on the anterior edge of each segment transversally towards the dorsal side of the trunk, connecting to the dorsal nerve cord (Additional File 6: Supplementary Figure 6; Additional File 7: Supplementary Figure 7b). During metamorphosis, the foregut will break from the larval tissue to connect with the brain and become the definite mouth of the juvenile (27). The patterns of innervation and gene expression remain very similar to that of the competent larvae (compare Figure 5 with Figure 7, and Additional File 3: Supplementary Figure 3 with Additional File 6: Supplementary Figure 6), except that now there are RGW-lir neurons on the lower mouth lip (lml; Figure 6g–h; Additional File 6: Supplementary Figure 6e–f). At this stage, *soxC* is broadly expressed in the mouth, and *six3/6* and *nk2.1* are expressed in the dorsal part of the foregut. *Otx* is now expressed in the boundary between the foregut and the midgut (Figure 7a–b, e–f, g–j), suggesting an additional role in the neural innervation of the foregut. Altogether, our findings indicate that significant changes in the neural architecture occur during metamorphosis, as the originally bilobed brain transforms into a ring and connects with the anterior part of the trunk, establishing the final nervous system architecture of the juvenile/adult.

**Figure 6.**
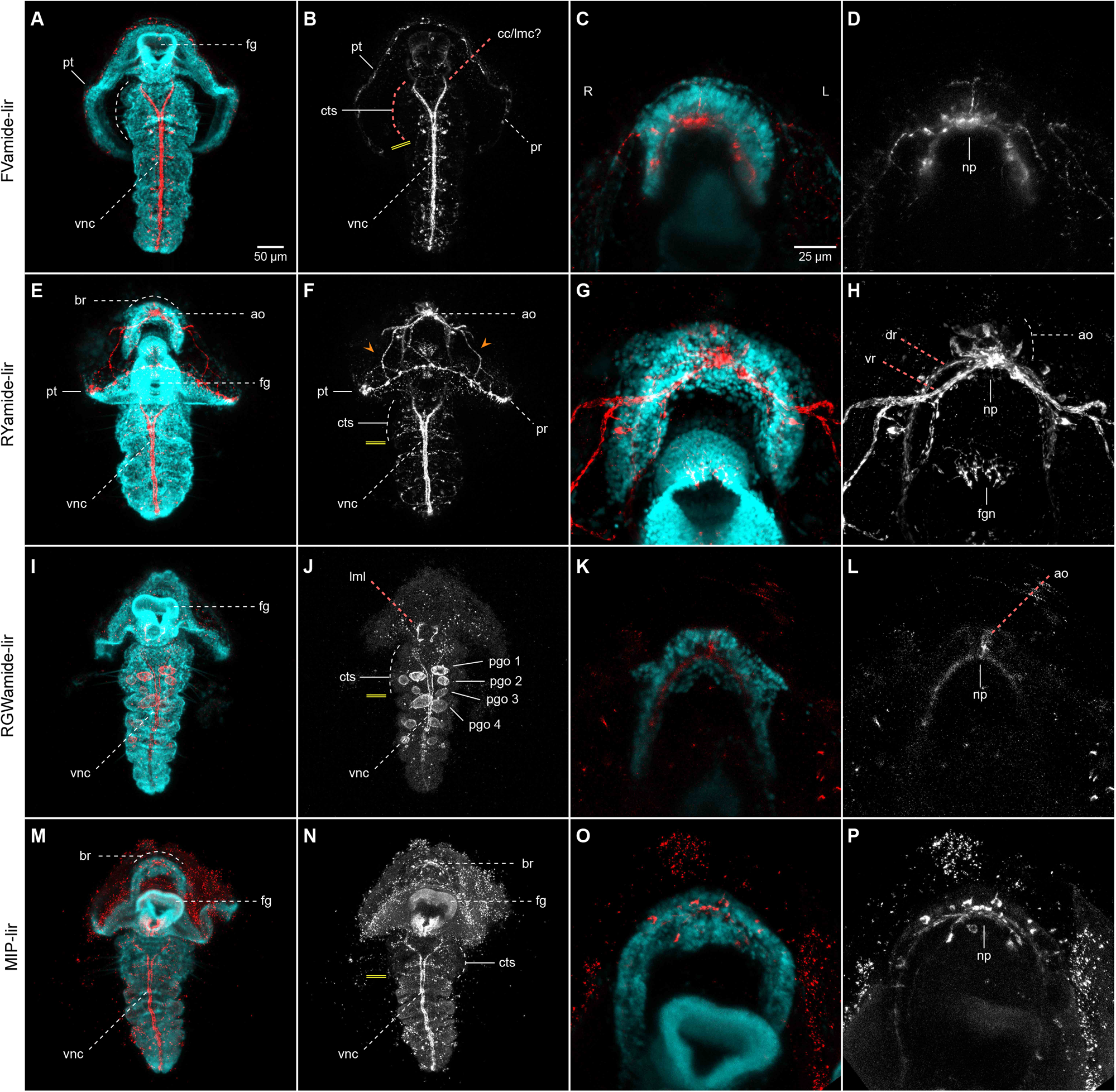
Neuropeptide-lir elements during metamorphosis. CLSM images of DAPI (cyan) and neuropeptide^+^ (red or white) elements during metamorphosis (∼ 3-4 wpf). Ventral views, with anterior to the top. **c**–**d**, **g**–**h**, **k**–**l**, **o**–**p** are close ups of the apical organ and brain in the same view as the respective larger image in **a**–**b**, **e**–**f**, **i**–**j**, **m**–**n**. **a**–**b**, **e**–**f**, **i**–**j**, **m**–**n** The brain connects with the ventral nerve cord (vnc), via circumesophageal connectives (lateral medullary cords (22) at the trunk thorax, made out of three ciliated thoracis segments (cts). Iterated **a–b** FVamide-lir, **e–f** RYamide-lir and **m–n** MIP-lir neurons and transverse lateral nerves are present in the segments of the trunk. **i–j** RWG labels the parapodial glandular organs (pgos). Double yellow line marks the division between thoracic and abdominal segments. ao: apical organ; ar: apical nerve ring; br: brain; cc: circumesophageal connectives; cts: ciliated thoracic segments; dr: dorsal root; fg: foregut; fgn: foregut nerve; lmc: lateral medullary cords; lml: lower mouth lip; np: brain neuropil; pgo: parapodial glandular organ 1**–** 4; pr: prototrochal ring; pt: prototroch; vnc: ventral nerve cord; vr: ventral root.

**Figure 7.**
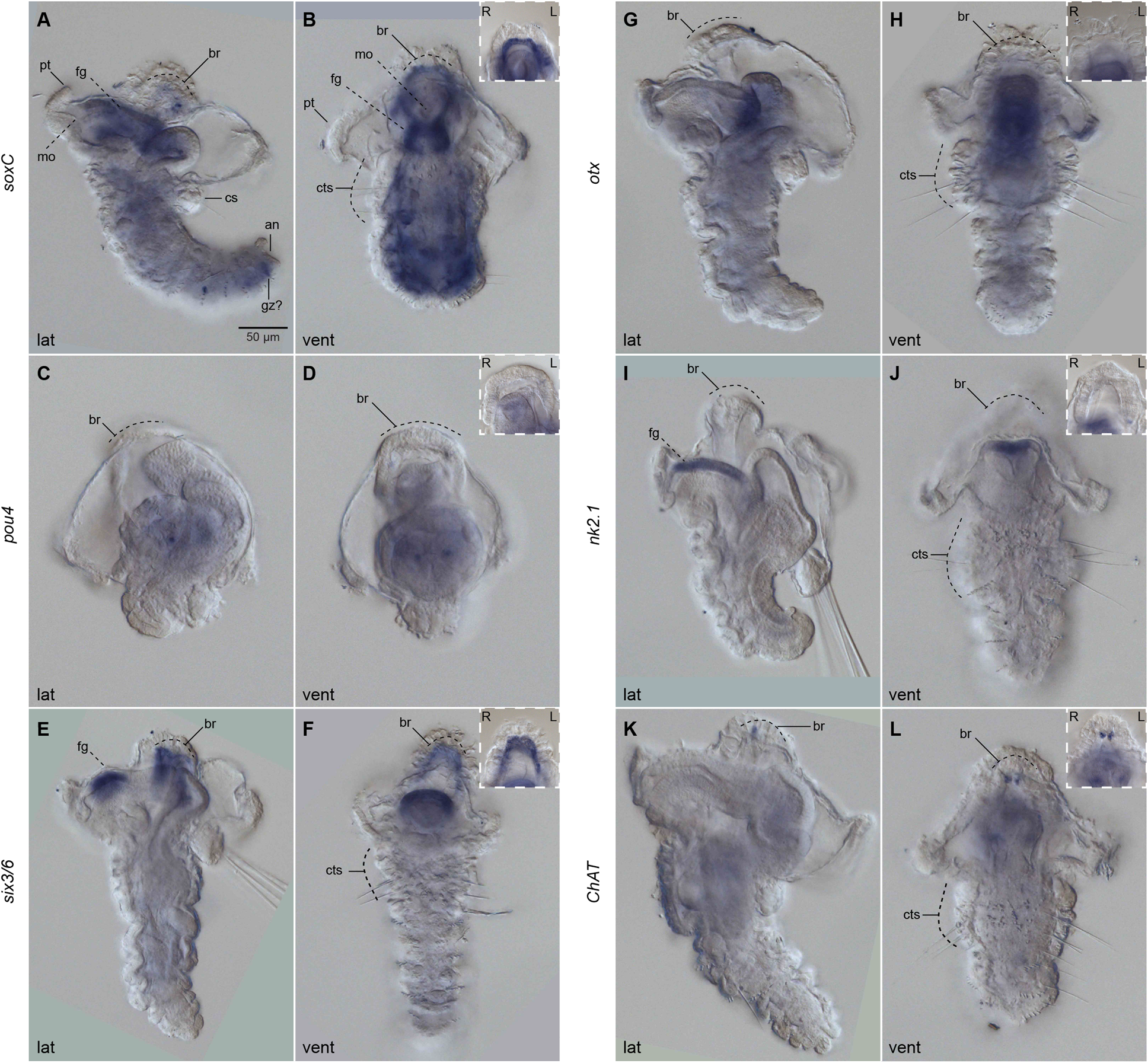
Neural development during metamorphosis. DIC images showing expression of *soxC*, *pou4*, *six3/6*, *otx*, *nk2.1* and *ChAt*. **a**, **c**, **e**, **g**, **i** Lateral views; **b**, **d**, **f**, **h**, **j**, ventral views. Insets are close ups focusing on the brain of the corresponding larger images. **a–b** *soxC*, **e–f** *six3/6* and **k–l** *ChAT* are expressed in the brain (br). **a–b** *soxC* is expressed throughout the trunk, the foregut (fg), and in the putative growth zone (gz). **e–f** *six3/6* and i**–j** are expressed on the dorsal side of the foregut, while **g–h** *otx* is expressed in the boundary between foregut and midgut. an: anus; br: brain; cc: circumesophageal connectives; cts: ciliated thoracic segments; dr: dorsal root; fg: foregut; fgn: foregut nerve; lmc: lateral medullary cords; lml: lower mouth lip; np: brain neuropil; pgo: parapodial glandular organ 1**–**4; pr: prototrochal ring; pt: prototroch; vnc: ventral nerve cord; vr: ventral root.

### The juvenile nervous system

After metamorphosis, the juvenile body subdivides into the head — with the fused prostomium and peristomium — and the trunk, further differentiating into three fused thoracic segments, seven abdominal segments, and the pygidium (27) (Figure 8). The mouth is anterior, and the brain ring is positioned dorsal to the roof of the foregut (23). The brain ring comprises 5HT-lir, FMRFamide-lir, and tubulin^+^ roots, connected via lateral medullary cords around the foregut to the vnc (9, 12, 23). The vnc has iterated 5HT-lir neurons in an otherwise continuous medullary cord with no breaks, as seen with *ChAt* expression (9, 12, 23). Consistently, FVamide-lir, RYamide-lir, RGWamide-lir, and MIP-lir localise to the ring-shaped brain that connects to the vnc with lateral medullary cords at the ciliated thoracic segments (Figure 8). FVamide-lir and RYamide-lir clusters of neurons (Figure 8a–d) and FVamide-lir, RYamide-lir, and tubulin^+^ peripheral nerves (Additional File 7: Supplementary Figure 7c–d) occur in the anterior part of each segment, with one tubulin^+^ pair of lateral nerves more prominent in each of the segments (ln; Additional File 7: Supplementary Figure 7c–d). Tubulin^+^ longitudinal nerve tracts run alongside the median vnc (cyan arrows; Additional File 7: Supplementary Figure 7c) and ventrolaterally (magenta arrows; Additional File 7: Supplementary Figure 7c–d). RGWamide-lir and MIP-lir nerves are also present in the mouth opening (Figure 8e–h). At this stage, *six3/6* and weakly *soxC* are expressed in the brain (Additional File 8: Supplementary Figure 8a–f). The latter is also expressed in the foregut and the putative posterior growth zone (gz), just before the pygidium (Additional File 8: Supplementary Figure 8a–c). Therefore, the definitive brain is primarily formed in the juvenile. However, the vnc neuroarchitecture is more elaborated at this stage than in the adult, as we describe below (12, 22).

**Figure 8.**
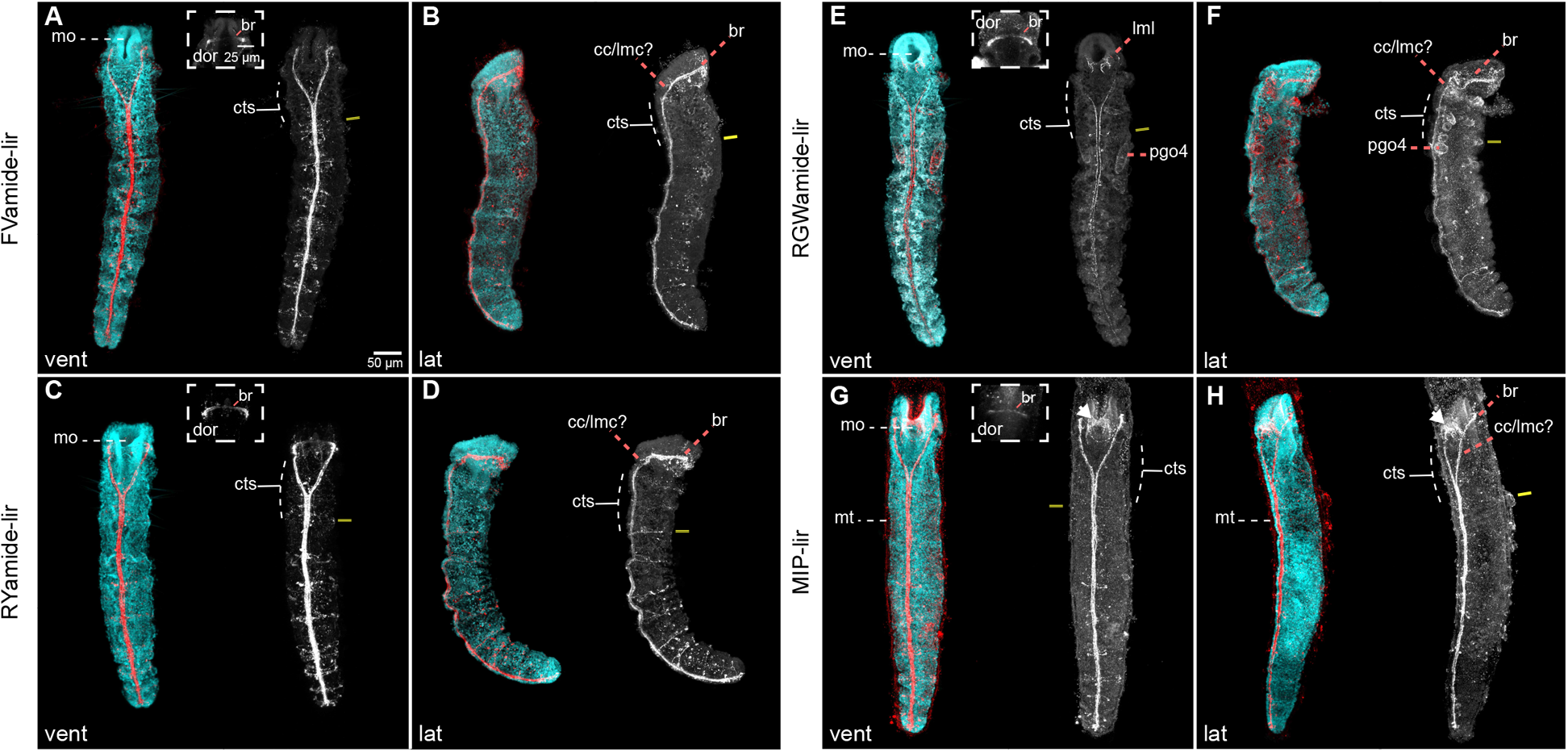
Neuropeptide-lir elements in the juveniles. CLSM images of DAPI (cyan) and neuropeptide-lir (red or white) elements in juveniles (>4 wpf). **a**, **c**, **e**, **g** Ventral views; **b**, **d**, **f**, **h** lateral views, with anterior to the top. The brain connects with the ventral nerve cord (vnc), via circumesophageal connectives (lateral medullary cords (22) at the trunk thorax, made out of three ciliated thoracic segments (cts). Iterated **a–b** FVamide-lir, **c–d** RYamide-lir and **g–h** MIP-lir neurons and lateral transverse nerves (ln) are present in the segments of the trunk. **e–f** RWG labels the parapodial glandular organs (pgos). Double yellow line marks the division between thoracic and abdominal segments. br: brain; cc: circumesophageal connectives; cts: ciliated thoracic segments; lmc: lateral medullary cords; lml: lower mouth lip; mo: mouth; mt: mucous tube; pgo: parapodial glandular organ 1**–**4; vnc: ventral nerve cord.

### The anterior adult neural structures

The head of the adult *O. fusiformis* includes a crown of tentacles formed from the fused prostomium and peristomium and a pair of ventrolateral eyes (22, 36) (Figure 9a). The FVamide-lir, RYamide-lir, RGWamide-lir, and MIP-lir nervous system is preserved throughout the ring-shaped brain, medullary cords, and vnc as seen in the juvenile (Figure 9; Additional File 9: Supplementary Figure 9). The neuropile of the brain is composed of parallel bundles of axons transverse to the lateral medullary cords, with FVamide-lir, RYamide-lir, RGWamide-lir, and MIP-lir neurons on the anterior and posterior edges (Figure 9b, d, f, h; Additional File 9: Supplementary Figure 9c, f, i, l). The FVamide-lir and RYamide-lir neuropil is wider than the RGWamide-lir and MIP-lir. The RYamide-lir neurons of the neuropil partially distinguish the dorsal and ventral roots of the brain as two concentrated bundles of neurites parallel to one another, separated by a less dense portion of neurites (Additional File 9: Supplementary Figure 9f), suggesting some level of compartmentalisation in the apparently simple ring-shaped brain of this annelid. Finally, there are FVamide-lir, RYamide-lir, RGWamide-lir, and MIP-lir longitudinal head nerves lateral to the brain (Additional File 9: Supplementary Figure 9b, f, j, n) that project anteriorly to the tentacles (22), and posteriorly into the trunk.

**Figure 9.**
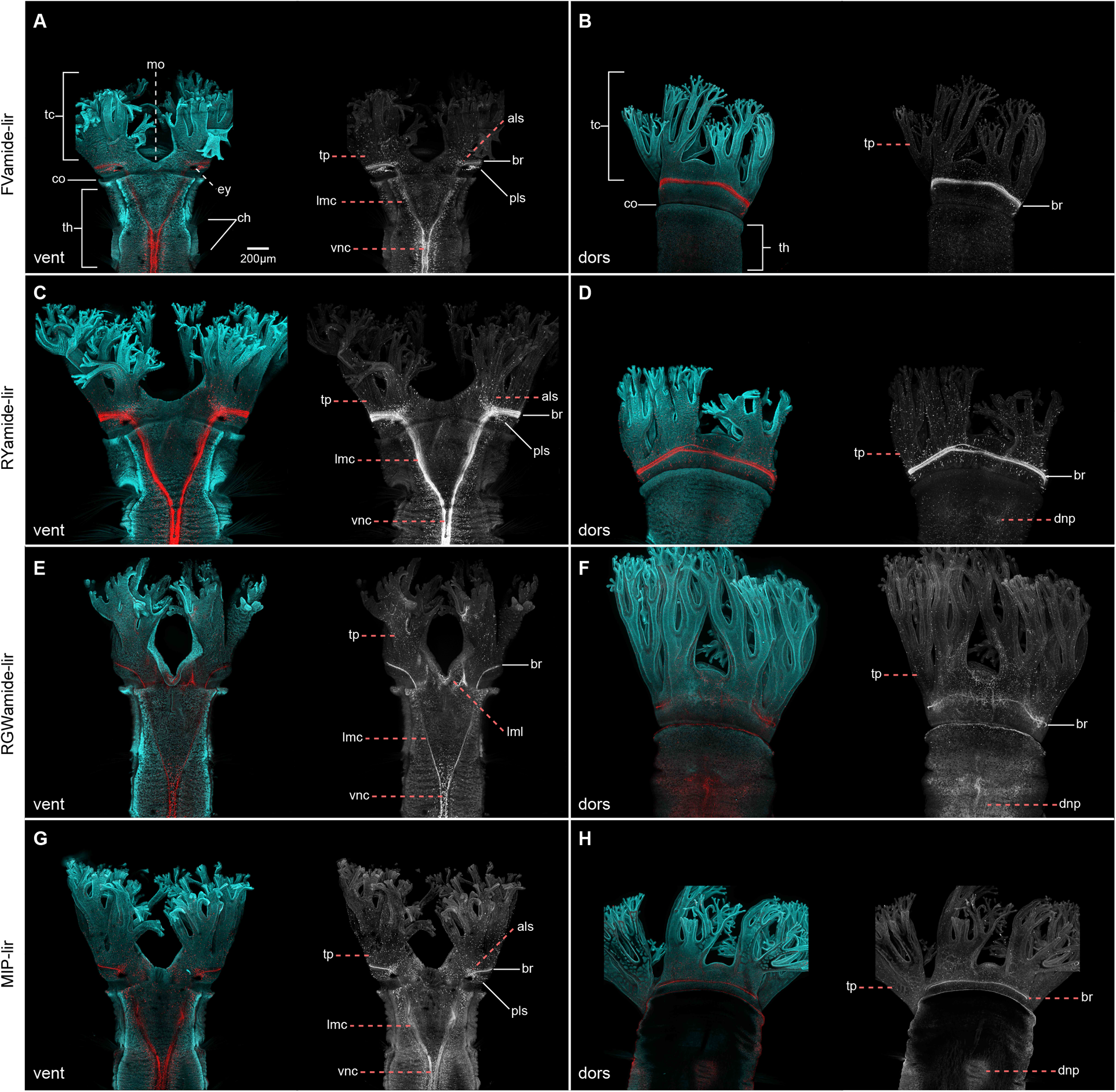
Neuropeptide-lir elements in the head of adults. CLSM images of DAPI (cyan) and neuropeptide-lir (red or white) elements. **a, c, e, g** ventral views; **b, d, f, h** dorsal views. **a, c, e, g** The FVamide-lir, RYamide-lir, RGWamide-lir and MIP-lir brain ring (br) is connected via lateral medullary cords (lmc) to the ventral nerve cord (vnc) at the position of the thorax (th). Each tentacle of the head contains a basiepidermal nerve plexus (tp), which projects from the brain. **b, d, f, h** Posterior to the head there is a dorsal nerve plexus (dnp). Surrounding each eye are clusters of somata oriented in an anterior-lateral (als) and posterior-lateral position (pls) position, showing FVamide-lir, RYamide-lir, and MIP-lir. als: anterio-lateral somata; br: brain; ch: chaetae; co: collar; dorsal nerve plexus: dnp; ey: eye; lmc: lateral medullary cord; lml: lower mouth lip; pls: posterior-lateral somata; tc: tentacle crown; th: thorax; tp: tentacle plexus; vnc: ventral nerve cord.

In addition to the brain, FVamide-lir, RYamide-lir, RGWamide-lir, and MIP-lir somata are present throughout the head tentacles (Figure 9 a–h) and surrounding the eyes (Figure 9a, c, e, g; Additional File 9: Supplementary Figure 9a, d, g, j). In these visual organs, a posterolateral cluster of neurons exhibits primarily FVamide-lir but also some RYamide-lir and MIP-lir signal, while RYamide-lir dominates in a second anterior cluster, which also shows some FVamide-lir and MIP-lir (Figure 9a, c, g; Additional File 9: Supplementary Figure 9a, d, j). However, this immunoreactivity is not part of the eye structure (21). A dorsal nerve cord composed of FVamide-lir, RYamide-lir, RGWamide-lir, and MIP-lir neurites and somata extends across the dorsal side of the body (Additional File 9: Supplementary Figure 9b, e, h, k). Some of these immunoreactivity patterns in the head support previously observed 5HT-lir and FMRFamide-lir clusters in other oweniids (43, 44). Our findings support that the adult brain and trunk nervous system are compartmentalised during the gradual reorganisation of the nervous system from larval and juvenile stages.

## Discussion

This study characterises the ontogeny of the nervous system in *O. fusiformis* from larvae to adulthood using a set of conserved cross-species antibodies and gene expression. The morphological landmarks presented here will serve as a foundation to understand larval development, metamorphosis, and post-larval morphogenesis in an annelid occupying a critical phylogenetic position, which will help to infer ancestral characters to Annelida and animals in general (Figure 10).

**Figure 10.**
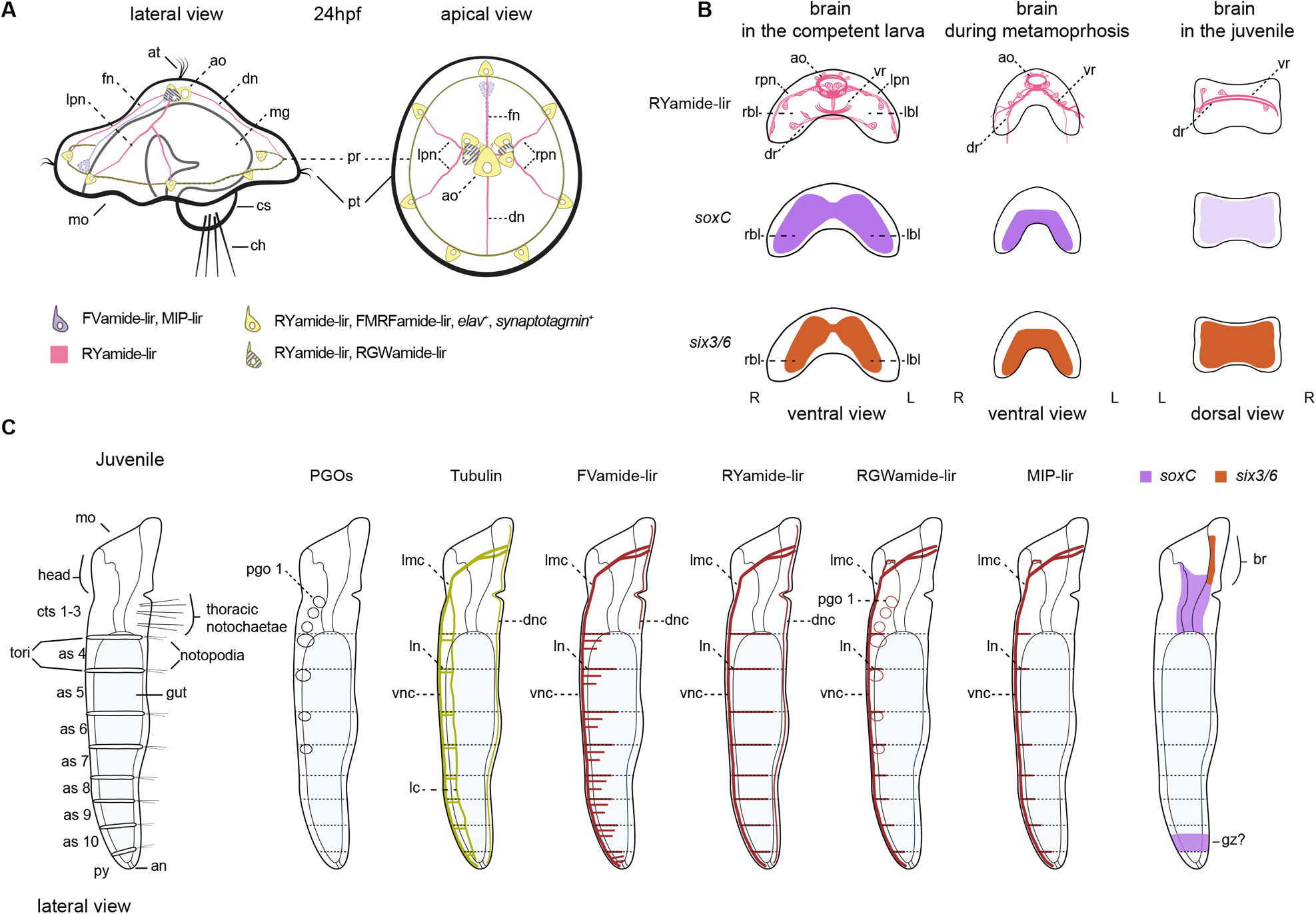
Diagram of neural development in *O. fusiformis*. **a** At 24 hpf there is an FVamide-lir, RYamide-lir, RGWamide-lir, MIP-lir and FMRFamide-lir apical organ with *elav*^+^ and *synaptotagmin*^+^ cells that connect to the prototroch ring (24). **b** The brain goes from a bilobed brain in the pre-competent larvae, to a ring in the juvenile **c** Pattern of immunoreactivity and *soxC* and *six3/6* expression in the juvenile. ao: apical organ; as: abdominal segment; at: apical tuft; br: brain; ch: chaetae; cs: chaetal sac; cts: ciliated thoracic segment; dn: dorsal nerve; dnc: dorsal nerve cord; dr: dorsal root; fn: frontarl nerve; gz: growth zone; lc: lateral cord; lmc: lateral medullary cords; lpn: left peripheral nerve; mg: midgut; pgo: parapodial glandular organ; pr: prototroch ring; pt: prototroch; rbl: right brain lobe; rpn: right peripheral nerve; vr: ventral root.

### The nervous system in the early larva

The mitraria larva largely derives from anterior/head tissues (29), and posterior territories are limited to a ventral epithelial invagination that will form the juvenile rudiment trunk (24, 27) and a small dorsal posterior tissue that includes the anus and chaetal sac (34). The larval neural system — composed of the apical organ and apical tuft connected to a prototroch ring — starts developing by 13 hours post fertilisation (hpf) and connects to the FMRFamide-lir prototroch ring by 24 hpf. The nervous system also includes seven FMRFamide-lir, *elav*^+^ and *synaptotagmin*^+^ neurons in the prototroch (24). Our findings support this early neural architecture of the mitraria larva and reveal further complexity and refinement, particularly in the apical organ and its connections to the prototrochal neural ring. As in *P. dumerilii*, the apical organ contains FVamide-lir, RYamide-lir, RGWamide-lir, and MIP-lir neurons in *O. fusiformis*, some of which are monociliated. All these neuropeptides form a neurosecretory centre that regulates the swimming behaviour of the larvae of *P. dumerilii* (31, 32, 45, 46). They are also present in the anterior neural systems of other annelid and spiralian larvae, as in *C. teleta*, and even directly developing species (30–32, 40). In *O. fusiformis*, the apical organ connects frontally, bilaterally, and dorsally to the prototroch (Figure 10a). The monociliary nature of the neuropeptide-lir neurons in the apical organ and the seven RYamide-lir neurons in the prototroch indicate they might have a sensory function (Figure 10a). They presumably integrate stimuli from the apical organ and the prototroch to control the shape of the episphere and the ciliary beating, thus influencing the locomotion and behaviour of the larva, without the need for excess neural wiring as hypothesised for larvae with monociliated cells (47).

The spatial patterns of immunoreactivity show notable similarities between *O. fusiformis* and *P. dumerilii*. In both larvae, RY (31), FV (30, 31, 48, 49), and MIP (32, 48–50) occur in ciliated sensory neurons. However, RY and RGW are expressed in interneurons that communicate to the synaptic nervous system in *P. dumerilii* (48, 49, 51, 52). Future studies of the connectome in *O. fusiformis* could clarify if this is true for *O. fusiformis*. Nonetheless, the presence of diverse neuropeptide sensory neurons, together with the deployment of staggered apical expression domains of transcription factors like *foxQ2*, *six3/6* and *otx* (11, 33, 53), support the evolutionary conservation of the apical region between annelids and spiralians and reveal anatomical traits of the anterior neural system of the ancestral “head swimming larva” of annelids.

### From a bilobed larval brain to an adult ring-shaped brain

With growth, the neural features present in the early larva become more elaborated (23, 24), and the adult nervous system develops, first with the condensation of nuclei that form the brain (Figure 10b) and later, with the patterning, elongation and subsequent evagination of the trunk. Nuclear staining, the expression of the anterior marker genes *ChAt*, *nk2.1*, *otx*, *pou4*, and *six3/6* (9, 33, 34), and neuropeptide immunoreactivity reveal that the pre-metamorphic larva has a bilobed brain (Figure 10b). This is consistent with classic morphological descriptions (27) and similar to the larvae of other “early branching” (54, 55) and more divergent annelids (15, 16, 56). The brain sits underneath a prominent neuropeptide-rich apical organ (Figure 10b), which comprises an apical ring and several neurons surrounding the monociliated apical tuft. Anterior and posterior FMRFamide-lir and 5HT-lir (23) and RYamide-lir axonal roots form a neuropil underneath the brain referred to as ventral and dorsal roots in other annelids, respectively (14, 41). Remarkably, this organisation changes with metamorphosis, as the bilobed brain forms a continuous *soxC*+ and *six3/6*+ band that compresses anteroposteriorly, bringing the dorsoventral roots closer to each other (Figure 10b). This results in the fusion of the brain lobes and roots into a double ring that forms the brain in the juvenile (23) and adult (22). While our data support a reorganisation of the brain from larval to adult stages (22, 25, 26), we were unable to determine the fate of the larval apical organ, and it remains unclear whether it integrates into the juvenile brain or is resorbed during metamorphosis with the apical tuft and prototroch.

From metamorphosis onwards, the roots of the brain neuropil connect with lateral medullary cords, ending into a medullary non-ganglionated, medially-condensed vnc in the trunk (12, 22). The presence of bundles of axons with distinct neuropeptide immunoreactivity in the adult brain ring suggests an unexpected level of compartmentalisation in this previously regarded “simple” brain (22) that might indicate the retention of the anterior and posterior roots (“ventral” and “dorsal”, respectively, according to traditional anatomical descriptions (14, 41)) seen in the larval and metamorphic stages in adult stages. This would challenge hypotheses based on the analysis of other oweniids that their ring-shaped brain is homologous to the dorsal (posterior) root neuropil of other annelids (43, 44). Despite its presumable compartmentalisation, there are no distinct ganglionic centres in the adult brain of *O. fusiformis*, unlike in more active annelids that exhibit structures like the mushroom bodies and nuchal organs (57, 58). Therefore, the brains of *O. fusiformis* and other representatives of the “early branching” clades gradually reorganise their morphology while retaining neuronal diversity during metamorphosis to form a continuous medullary cord with the vnc, perhaps associated with a transition to a more sedentary, tube-dwelling lifestyle as adults.

### From a juvenile rudiment to the trunk nervous system

The trunk of oweniids forms as an invagination of the ventral epithelium of the larva (27, 28) with the deployment of conserved anterior-posterior and trunk-patterning programmes like the *hox* genes (29). While neurogenesis, as revealed by the expression of *elav, synaptotagmin* (24), and *soxC* (this study), is predominant in the apical organ and brain region in the early larva, it mainly occurs in the developing trunk before metamorphosis. As in other annelids (9, 12), the trunk nervous system develops as a paired medially-condensed vnc, but, most notably, it also includes a single dorsal nerve cord connected to the ventral one by segmentally iterated lateral nerves. During metamorphosis, additional ventrolateral longitudinal cords form, giving the trunk nervous system an orthogonal appearance that has been hypothesised to be the ancestral pattern for annelids (59) and other spiralians, such as flatworms and nemerteans (60, 61). A ganglionated ladder-like vnc thus likely evolved independently multiple times in annelids and animals (9, 12). As the juvenile worm matures into adulthood, more neurons appear along the vnc, resulting in a continuous medullary cord with no apparent breaks (12, 22, 43). However, the lack of segmented ganglia in the vnc of *O. fusiformis* does not exclude the presence of clusters of 5HT-lir (9, 12, 23) and FVamide-lir, RYamide-lir and MIP-lir (this study) neurons in each segment. Parapodial glandular organs (PGOs) (42) develop in each of the first seven segments (27) and show RGWamide-lir, which combined with the cilia of the thoracic segments and the neuropeptide-lir and tubulin^+^ lateral nerves of the abdominal segments, define positional landmarks along the anterior-posterior axis that would aid in the study of trunk formation in *O. fusiformis* (Figure 10c). Concurrent with the maturation of the brain and trunk nervous system, the immunoreactivity in the larval foregut and definitive oesophagus changes. In *O. fusiformis,* the foregut of the competent larvae is innervated by 5HT-lir and FMRFamide-lir (23, 24), and FVamide-lir and RYamide-lir neurons and axons (this study); and by 5HT-lir (23) and RGWamide-lir and MIP-lir (this study) in the juvenile stage. FMRFamide-lir neurons and axons innervate the enteric nervous system of juvenile annelids like *C. teleta* (15). At the same time, MIP is also present in the stomatogastric nervous system in dinophilids (40), and it plays a role in the feeding behaviour of *P. dumerilii* larva (50), suggesting a conserved neuropeptide-mediated control of feeding in annelids.

## Conclusions

Our study describes the transition of the nervous system from the early larva to the adult stage in the annelid *O. fusiformis*, a representative of Oweniidae and the sister lineage to all remaining annelids. The initial larval neural system comprises an apical organ connected to a prototrochal ring and the chaetal sac through several neurites. Soon, a bilobed brain forms underneath the apical organ, connecting with other larval tissues and the developing juvenile trunk in its anterior part. During metamorphosis, the lobes, and the ventral and dorsal roots fuse to form a ring-shaped brain, following a similar trend of reorganisation of the neural architecture as in other “early branching” annelids like magelonids and chaetopterids (22, 25, 26). However, our findings indicate that the larval and adult nervous systems are not as simple as previously thought in *O. fusiformis* and retain similarities with more deeply nested annelids, particularly at the larval stages. Future studies of the detailed connectome of the mitraria larva will help to understand how these anatomical similarities translate into conservation of behaviours and physiological functions, illuminating how neuropeptidergic systems might have contributed to the evolution of biphasic life cycles.

## List of abbreviations

als: anterio-lateral somata
an: anus
ao: apical organ
ar: apical nerve ring
at: apical tuft
br: brain
CLSM: confocal laser scanning microscopy
cc: circumesophageal connective
chn: chaetal sac nerve
co: collar
cs: chaetal sac
cts: ciliated thoracic segment
dn: dorsal nerve
dnc: dorsal nerve cord
dr: dorsal root
fg: foregut
fgn: foregut nerve
fn: frontal nerve
gz: growth zone
jr: juvenile rudiment
lc: lateral cord
lmc: lateral medullary cord
lml: lower mouth lip
ln: lateral transverse nerve
lpn: left peripheral nerve
mg: midgut
MIP: myoinhibitory peptide
mo: mouth
mt: mucous tube
ne: neurite
np: brain neuropil
nph: nephridia
pgo: parapodial glandular organ
pls: posterior-lateral somata
pr: prototrochal ring
pt: prototroch
so: somata
rg: refringet globule
rpn: right peripheral nerve
tc: tentacle crown
th: thorax
tp: tentacle plexus
vnc: ventral nerve cord
vr: ventral root

## Declarations

### Ethics approval and consent to participate

Not applicable.

### Consent for publication

Not applicable.

### Availability of data and materials

The datasets used and analysed during the current study are available from the corresponding author upon reasonable request.

## Authors’ contributions

AMCB and JMMD designed the study. AMCB and RD performed all the immunostainings and fluorescence imaging. AMCB performed the expression analyses and imaging of gene expression. EAW and GJ contributed with reagents and sequencing data. AMCB, RD and JMMD built the figures. AMCB drafted the manuscript. All authors contributed to data interpretation and manuscript writing.

## Acknowledgements

We thank all members of the Martín-Durán lab for their support. We thank Océane Seudre for amplifying and synthesising the *soxC* probe and Paul Kalke for recommendations regarding immunostaining of the adult heads. We also thank the Station Biologique de Roscoff for their help with collections and animal supplies.

---

Supplementary Figure 1 Alignment of the neuropeptide precursors *P. dumerilii* (30–32), *C. teleta* (62) and *Owenia fusiformis* (29). Representative mature peptides and conserved dipeptides are highlighted in red and bold, respectively.

Supplementary Figure 2 MIP-lir elements in the 24hpf mitraria. MIP-lir cells include several cells as part of the apical organ (ao) and one cell anterior to the foregut (white arrow), including a MIP-lir frontal nerve (fn). Inset in **b** is a close up of the apical organ (ao) in the same view as the larger image. ao: apical organ; at: apical tuft; cs: chaetal sac; fn: frontal nerve; mo: mouth.

Supplementary Figure 3. Neuropeptide-lir elements in the competent larvae. CLSM images of DAPI (cyan), acetylated tubulin (yellow) and neuropeptide-lir (red or white) elements in the competent larvae (∼ 3 wpf). Lateral views, with anterior to the left. **c**, **f**, **i**, **l** are close ups of the juvenile rudiment in the same view as the respective larger image in **b**, **e**, **h**, **k**. **a–c** FVamide-lir cells and MIP-lir cells in the apical organ connect via FVamide-lir and MIP-lir circumesophageal connectives (cc) to the ventral nerve cord (vnc) of the juvenile trunk rudiment (jr), and via **a–b** FVamide-lir, **d–e** RYamide-lir and **j–k** MIP-lir frontal (fn), dorsal (dn) and peripheral nerves (closed orange arrow heads) to the **a–c** FVamide-lir, **d–f** RYamide-lir and **j–l** MIP-lir prototrochal ring (pr). See also Figure 2. **d–f** RYamide-lir and **j– l** MIP-lir peripheral nerves also branch out to the chaetal nerve (chn) (open pink arrowheads). The foregut is innervated by **a–c** FVamide-lir and **d–f** RYamide-lir cells and neurites. By this stage the juvenile rudiment has a vnc and a **a–c** FVamide-lir and **d–f** RYamide-lir dorsal nerve cord (dnc). **g–i** RGWamide-lir cells are only present in the apical organ. **j–l** MIP-lir is present in the anterior part of the foregut (white arrow). an: anus; ao: apical organ; at: apical tuft; br: brain; cc: circumesophageal connectives; chn: chaetal sac nerve; cs: chaetal sac; dn: dorsal nerve; dnc: dorsal nerve cord; fg: foregut; fgn: foregut nerve; fn: frontal nerve; jr: juvenile rudiment; mg: midgut; mo: mouth; pr: prototrochal ring; pt: prototroch; vnc: ventral nerve cord.

Supplementary Figure 4 Tubulin^+^ elements in the competent. CLSM images of beta-tubulin (**a–e**) and alpha-acetylated tubulin (**f–j**) in the competent larvae (∼3 wpf). **a–c**, **g–h** apical views; **d–e**, **i–j**, lateral views; **f** ventral view. **a–c**, **g–h** the apical organ (ao), associated with an apical tuft (at) and apical nerve ring (ar) is positioned above the brain (br). Ventral (vr) and dorsal (dr) roots **c**, **h** make the neuropil of the brain, that connects with the **d** cirucomesophageal connectives (cc), and ultimately with the ventral nerve cord (vnc) **d–e**. **f**, **i–j** Tubulin^+^ peripheral nerves (fn, dn, and orange arrowheads) connect the apical organ with the prototroch ring (pr). ao: apical organ; an: anus; ar: apical nerve ring; at: apical tuft; br: brain; cb: chaetoblast; cc: circumesophageal connectives; chn: chaetal sac nerve; cs: chaetal sac; dn: dorsal nerve; dnc: dorsal nerve cord; dr: dorsal root; fg: foregut; fgn: foregut nerve; fn: frontal nerve; jr: juvenile rudiment; mg: midgut; mo: mouth; nph: nephridia; pr: prototrochal ring; pt: prototroch; vnc: ventral nerve cord; vr: ventral root.

Supplementary Figure 5 SoxC orthology and early mRNA expression. **a** Maximum likelihood orthology assignments of *soxC*. **b** DIC images showing expression of *soxC* during gastrulation (9hpf) and early mitraria (24hpf). Asterisks mark the animal/apical pole an: anus; bp: blastopore; cs: chaetal sac; fg: foregut; mo: mouth; pt: prototroch.

Supplementary Figure 6 Neuropeptide-lir elements during metamorphosis. CLSM images of DAPI (cyan) and neuropeptide-lir (red or white) elements during metamorphosis (∼ 3 4pf). Lateral views, with anterior to the top. **a–h**The brain connects with the ventral nerve cord (vnc), via circumesophageal connectives (lateral medullary cords (22) at the trunk thorax, made out of three ciliated thoracis segments (cts). The foregut (fg) has **b** FVamide-lir, **f** RYamide-lir and **h** MIP-lir neurons and cells. **e–f** RWGamide labels the parapodial glandular organs (pgos), and the lower mouth lip (lml). Double yellow line marks the division between thoracic and abdominal segments. ao: apical organ; an: anus; br: brain; cc: circumesophageal connectives; cts: ciliated thoracic segments; dn: dorsal nerve; dr: dorsal root; fg: foregut; fgn: foregut nerve; lmc: lateral medullary cords; lml: lower mouth lip; np: brain neuropil; pgo: parapodial glandular organ 1**–**4; pr: prototrochal ring; pt: prototroch; vnc: ventral nerve cord; vr: ventral root.

Supplementary Figure 7 Tubulin^+^ elements during metamorphosis and juvenile. CLSM images of acetylated tubulin. **a–b** Larvae undergoing metamorphosis. **c–d** >4 wfp juvenile. **a–b** Tub^+^ peripheral nerves (orange arrowheads) in the remaining episphere of the larva keep connecting the brain to the prototrochal ring (pr). **a–d** The brain connects with the ventral nerve cord (vnc), via circumesophageal connectives (lateral medullary cords (22) at the trunk thorax, made out of three ciliated thoracis segments (cts). The vnc is composed of two robust longintudinal tracts, and two more lateral tracts (magenta arrows). On the anterior border of each segment, there is a pair of lateral transverse nerves (ln) that connect to lateral ventral-lateral longitudinal cords (magenta arrows). Double yellow line marks the division between thoracic and abdominal segments. ao: apical organ; br: brain; cc: circumesophageal connectives; cts: ciliated thoracic segments; dr: dorsal root; fn: frontarl nerve; lmc: lateral medullary cords; ln: lateral transverse nerves; nph: nephridia; pr: prototrochal ring; pt: prototroch; vnc: ventral nerve cord; vr: ventral root.

Supplementary Figure 8 Neural development in juveniles. DIC images showing expression of *soxC*, *pou4*, *six3/6* and *otx*. **a**, **d**, **g**, **j** Lateral views; **b**, **e**, **h**, **k** ventral views; **c**, **f**, **i**, **l** dorsal views. **a–c** *soxC* and **d–f** *six3/6* are expressed in the brain (br). **g–i** *pou 4* and **j–l** *otx* have no longer any neural expression. **a–c** *soxC* is expressed in the foregut (fg), and in the putative growth zone (gz). br: brain; fg: foregut; gz: growth zone; mo: mouth.

Supplementary Figure 9 Neuropeptide-lir elements in the adults. CLSM images of neuropeptide-lir close ups of images in Figure 9. **a–b, e–f, i–j, m–n** ventral views; **c–d, g–h, k–l**, **o–p** dorsal views. **a, e, i, m** Views of the eye showing FVamide-lir, RYamide-lir and MIP-lir, antero-lateral (als) and postero-lateral (pls) somata. **b**, **f**, **j**, **n** Lateral head neurites (lhn) extend toward the tentacles and the trunk. **c**, **g**, **k**, **o** Longitudinal dorsal nerve cord (dnc). **d**, **h**, **l**, **p** Brain ring with associated neurites (ne) and somata (so). als: anterior-lateral somata; br: brain; dnc: dorsal nerve cord; ey: eye; lhn: lateral head neurites; lmc: lateral medullary cord; ne: neurite; pls: posterior-lateral somata; so: somata; tp: tentacle plexus.

