## Additional File 1: Supplementary Figure 1 for "The development of the adult nervous system in the annelid *Owenia fusiformis*"

**FV**

```
P. dumerilii FVamide precursor AEE25642.1    M------SVG GWPLLL---- ---------- --LIIAHGS- SLQAQDADSE NMRQLLQSMM LEDALSANEN SLGG------ --QNAPGYI GDDKRAAHRF VGKRDS----
C. teleta FVamide precursor ELU03112.1       C------E-G GKAEDI---- ---------- --IQNAYGEL SL----GELR NLEDYLAEYY FNSLIYSNDA AVDDYSYLLS SQQGSVPGQ- PQKREIDERF YQLRRAPPHS
O. fusiformis FVamide precursor CAH1786749.1 MLSDLNQSPG GETEDISQVD GYRMDLQVDL VPLVSCLFAL VITASGQSSN KLGLSSASVC RQSELWNSAC SLCEQLDLGF TQEGCC--LE TEYQAVCHAC LSNDDCLLTG

P. dumerilii FVamide precursor AEE25642.1    ---------- -EKRRLFVGK REDDEDEEKR AHMFVGK--- REDDKRAHRF V--------- --------GK REEEKRRLFV GKRD------ -----D-DKR NRMFVGKRSD
C. teleta FVamide precursor ELU03112.1       SYPRGYYFTM RQRRPQFVGK RALDSFTDSL ALMSASEFSQ PQSNNILHTG V--------- ---------K RAAPN---FI GKRD------ -----GLEKR AHMYIGRRSP
O. fusiformis FVamide precursor CAH1786749.1 LVEKRRNMFV GKRRNMFVGK RR-NMFVGKR RNMFVGKRSY YEEDKMQPEL VRKPVIDELD RLTAEILASK SMAKRQRMFV GKRDLNGENI ANEIEELEKR RNMFVGKRDP

P. dumerilii FVamide precursor AEE25642.1    FDDNDFEYDE KRRYFIGKRD GISRDIRRLW IGKRGDGYDY KRPHNFVGKR GHMFVGKR-- --DDFDLAED KR-RMFIGKR MYDDLEDYD- DKRARMFIGK RAHMFVGKRY
C. teleta FVamide precursor ELU03112.1       M--SDLTLDD ----FLD--D TYSED----- --------QS KRAPMFVGKR NRMFVGKRFD DEDDFLSSEE KRAHLFVGKR EREDMELQEL AKLLEMESSK RAPAFVGKR-
O. fusiformis FVamide precursor CAH1786749.1 IDE-----EK RRNMFVGKRE SIDEE----- ---------- KRRNMFVGKR DQM------- -------EEQ KRRNMFVGKR ESIDEE---- ---------K RRNMFVGKR-

P. dumerilii FVamide precursor AEE25642.1    MDDLDDEESK R-RMFIGKR- ------NR-M FVGKRDDELD QDALSLDKRR QLYIGKKFAV EPAEETRDLE EGEEKRNRMF IGKRSAEVRQ PADASVSSQ- -----
C. teleta FVamide precursor ELU03112.1       -DSYENEEDK RAHTFVGKK- ------MAPS FIGKR----- --------RA PGFIGRRQA- -PGFVG---- ----KRNSFN IVKRSATEDS NNSDLQDN*- -----
O. fusiformis FVamide precursor CAH1786749.1 -ESIDEE--K RRNMFVGKRD QIEEQKRRNM FVGKRE---- --SIDEEKRR NMFVGKRESI ---------- -AEEKRRNMF VGERNPLDLL SNDDEVEKRA PKFVG
```

**RY**

```
P. dumerilii RYamide precursor AEE25645.1    M-S------- ---------- IDSRTLILA- LAVFVTLAFA ---------- ----ED--KD DQSTSVAADE DDKRGTLL-- ---------- -------RYG
C. teleta RYamide precursor ELT93665.1       MQMSIINLIT TIKKQESPIE MDMWTILLS- MVVFVGV--- ---------- VSCSEENQIN KRDVSDVAEG DNT------- ---------- -----FDREA
O. fusiformis RYamide precursor CAH1772730.1 M-LGVLSDST EVKNKNIRRL TDYRTKMMGW MIAFTTLGLV VIVTAQPTNG IQNIDDDISD NDDLELLLDD TDPLVNKLIQ LFLDHEQTYD TDDDPFQRSI

P. dumerilii RYamide precursor AEE25645.1    RGSLMRYG-R GSLMRYGKRG TLMRYGKRDD EEFYDDDDVD AIKRVFRYGK R---GDDDYK RLFRWGKRSD DYVPEDG-LD EDKRGTLMRY GKRVFRYGKR
C. teleta RYamide precursor ELT93665.1       RAAIFRYGKR VPIFRYGKR- -IFRYGKRAD QEV-EDSDMD LEKRLFRYG- RSGSMPTTYK RLFRYGKRAE DDVEEDDEIE AVKR--LFRY GKR-------
O. fusiformis RYamide precursor CAH1772730.1 RGNLLRYGKR GALFRYGKRQ SFMRYGKRGG ---------- ----LFRFGK RASSGSTAGK RIFRWGKRSE ETA------- -EKRGNLMRW GKRA------

P. dumerilii RYamide precursor AEE25645.1    SDDDSASEDM AKRVFRYGKR NSVDLDDLIE EAKRRIFRYG RDIRAPQAPH VPFRFG---- -EE
C. teleta RYamide precursor ELT93665.1       GEINTRT--- ---------- ---------- ---------- ----AAQQPS VPFRFGE--- RE*
O. fusiformis RYamide precursor CAH1772730.1 GDSDFNTE-- ---------- NEI-ADDLSM RATSK--HVE RDDKL----H VPFRFGRDES EEQ
```

**RGW**

```
P. dumerilii RGWamide precursor AFS33094.1     MKLQGVVASW L---FFGLVI LE-------A TYAADSGTSQ IDKRRGWGKR DSEGEVDKRR GWGKRGALDP ELESEADKRR GWGKRS---S LEEMEKRRGW GKRTSVEEEK
C. teleta RGWamide precursor ELU14209.1        MLLNHLILLS LALAFIPAVS FA-------S DELVDEEASE MDKRRGWGKR -SDSEMDKRR GWGKR-SFDE DSE-EVAKRR GWGKRSEAIN LDEVEE---- ---SIAEMDK
O. fusiformis RGWamide precursor CAH1783991.1 MKL------Y IAIALLGLVV FELCTDFARA DDPAD--ATD MDKRRGWGKR ---------- GWGKR----- ---------- GWGKRSE--- -EQMDD---- ---DVLDEYV

P. dumerilii RGWamide precursor AFS33094.1     RRGWGKRDFE E--EEEMDKR RGWGKRSLEE AEEKRRGWGK RLTIDALEDG IDKRRGWGKR SQMEDEEEKR RGWGKRSWPK --DSAEA--C SEMRQAVVYY INSAIETEGM
C. teleta RGWamide precursor ELU14209.1        RRGWGKRSEL EAMQSEFAKR RGWGKRADDM EILKRRGWGK R-SEDANE-- MDKRRGWGKR SL---EAEKR RGWGKRS-PK IEEFSEFVDC QEVKEKIFYH VSLALQLDAQ
O. fusiformis RGWamide precursor CAH1783991.1 KRGWGKR--- ---------- -GWGKR---- ---------- ---------- -----GWGKR SSSIAES--- ---------- ---------C DEMNPDLVYL IYKAVEAEAQ

P. dumerilii RGWamide precursor AFS33094.1     RVKQCGGDPI VTP
C. teleta RGWamide precursor ELU14209.1        RANIC---PA EK*
O. fusiformis RGWamide precursor CAH1783991.1 RMNSCNK--- VDS
```

**MIP**

```
P. dumerilii MIP precursor AFV92893.1     MD--RVTITC FSLCLAS--V LIPLVHSEEN V------DLD ED---KRAWM KNNI------ ---------- ---------- -----AWGKR GWKQGASYSW GKRDSEGDGL
C. teleta MIP precursor ELU01469.1        MASCRLLLTV ITLVICSLVV LADDPETEEQ AQDLVPHDMD QDLMDKRKWG SNSMRVWGKR DGDDEME-MD GGAEKRKWGG NNNMRVWGKR KWGANSMRVW GKRSE-----
O. fusiformis MIP precursor CAH1786447.1 MAR-KELFSI FCILLIPCII HCENEKTDSN A--IQSTKTA ESRVDELKDT LEDLKTLAKE EDNDIISSVQ EDPAKRNWKS G---QFWGKR GDGWKSNQFW GKRDGD----

P. dumerilii MIP precursor AFV92893.1     MSDEEKRAWN -KN------- -NMRVWGKRS DEDD-----K RGWKDSSMRV WGKRAGEDDN NKRWGKNNL- --------RV WGKRAD---D LEVLEDKRAW GDNNMRVWGK
C. teleta MIP precursor ELU01469.1        LPEEEKRKWG GSN------- -TMRTWGKRA DDNEEDELAK RKWGSNSMRV WGKRA--DDN KRKWGSNSM- --------RV WGKRAD---D -DMDESKRGW KNNNMRVWGK
O. fusiformis MIP precursor CAH1786447.1 --DEDKRAWK SGQFWGKRAW KSGQFWGKRS DDDE-----K RAWKSGQF-- WGKRAGDELD KRNWKSGQFW GKRDWKSGQF WGKREDPAIE YEDNQDKRNW KSGQF--WGK

P. dumerilii MIP precursor AFV92893.1     RSD--LEDDK RAWNKNSMRV WGKRDMEED- ---------- -------ED- ----NKRAWK GQSARVWGKR --ADEDD--K RGWNGNSMRV WGKRG--WHG NGVRQWGKRL
C. teleta MIP precursor ELU01469.1        RADDEIDEDK RKWGSNSMRV WGKRSADDDA ELAAAVPHAI VKRSLDSEDF TDDMEKRRWG GNDMRVWGKR --RSRADGPK RSWKTNVMRV WGKRG--WAD NNMRVWGKRA
O. fusiformis MIP precursor CAH1786447.1 R---DMDEDK RKWKSGQF-- WGKRSVDSEN TNDVQVD--- -----DSIDS THSIAKRSTD WKSASLWGKR DWKSGQFWGK RAWQNPGS-- WGKRGDSWKA GNF--WGKRA

P. dumerilii MIP precursor AFV92893.1     HLDDVEPILD DEESKRAWAK NNMRVWGKRS --TDNV-RNM KAVVA----- -------EPA EVAADAE--- SA----KSS
C. teleta MIP precursor ELU01469.1        ---------D EGAEKRAWVG DKSLSWGKRS --DNEVIRNL LAEQE----- -------KEN ELREYLG--- SRLYEDEV*
O. fusiformis MIP precursor CAH1786447.1 D--------- ------SWKS GNF--WGKRD AGEDDVLDTL ITERVMKYFN NDSNDCLDRK ELQQFINWLT TLGMADMEL
```
