## Supplementary figures and images for "The development of the adult nervous system in the annelid *Owenia fusiformis*"

### Additional File 2: Supplementary Figure 2

#### Additional File 2: Supplementary Figure 2

MIP-lir

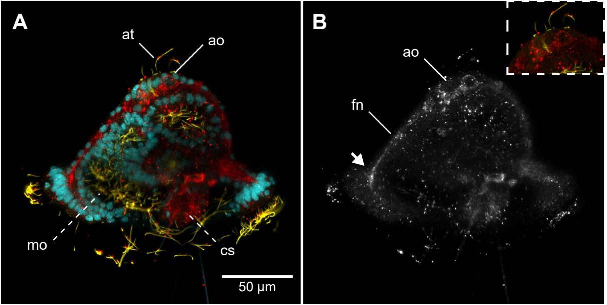

### Additional File 3: Supplementary Figure 3

##### Additionial File 3: Supplementary Figure 3

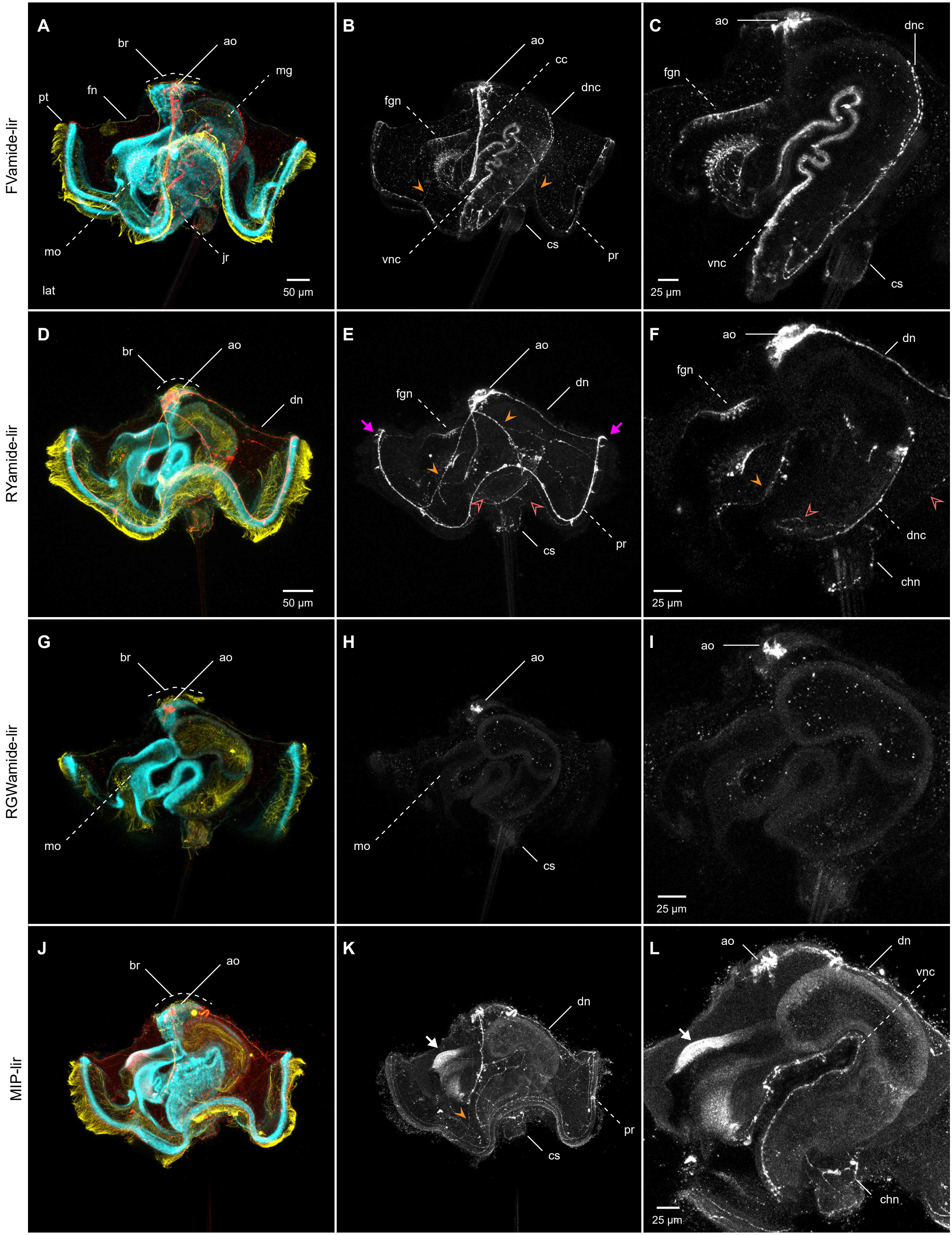

### Additional File 4: Supplementary Figure 4

#### Additional File 4: Supplementary Figure 4

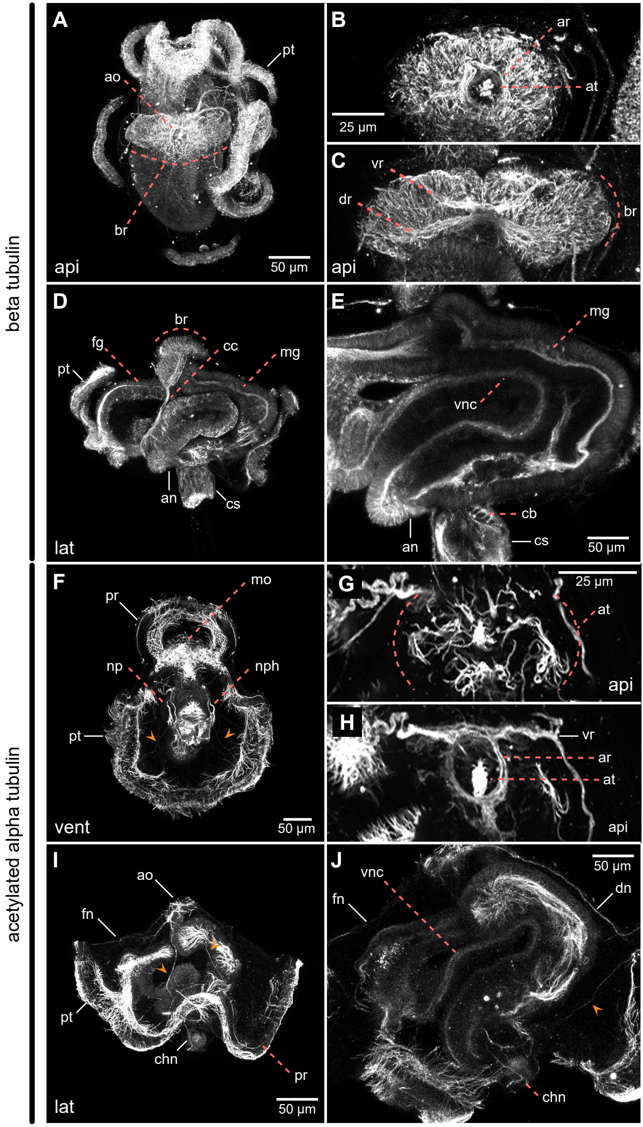

### Additional File 5: Supplementary Figure 5

#### Additional File 5: Supplementary Figure 5

**A**

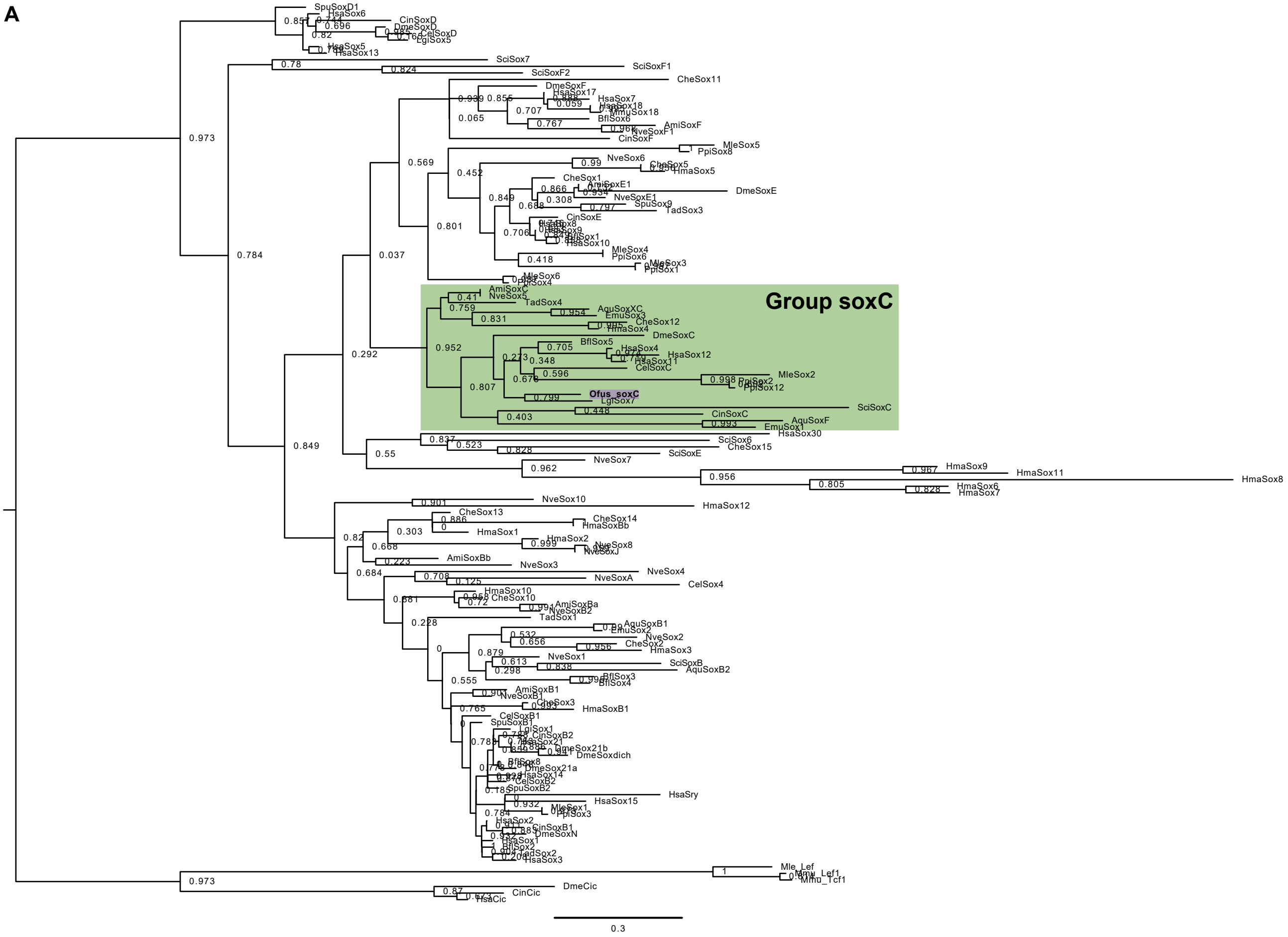


**B**

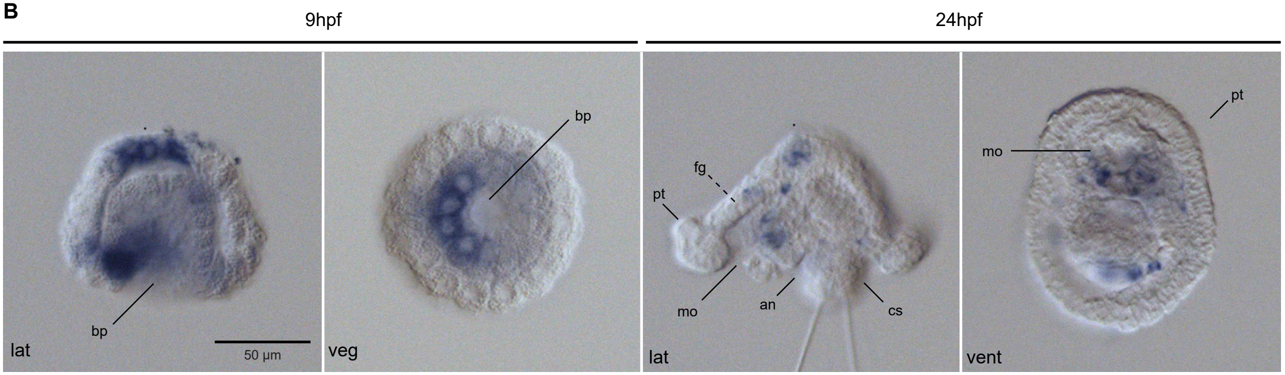

### Additional File 6: Supplementary Figure 6

#### Additional File 6: Supplementary Figure 6

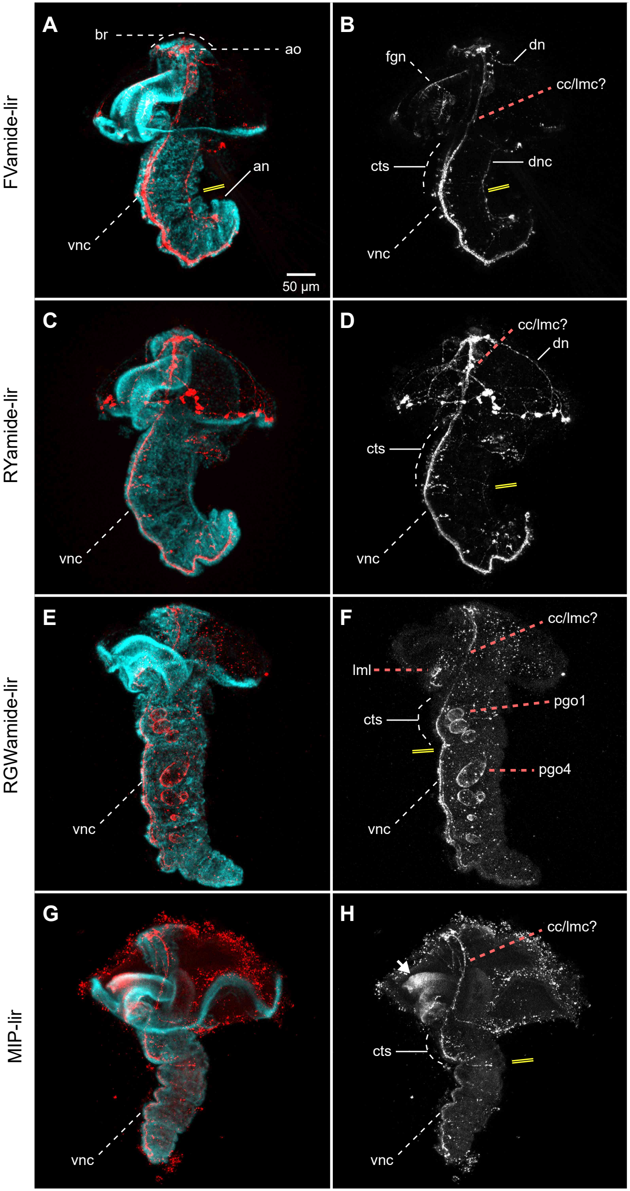

### Additional File 7: Supplementary Figure 7

#### Additional File 7: Supplementary Figure 7

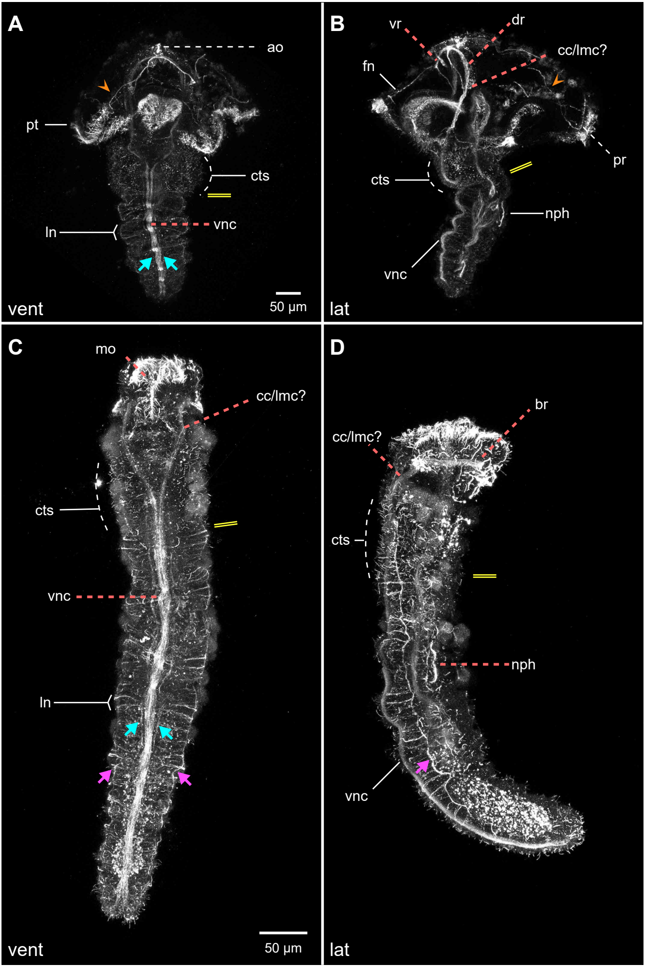

### Additional File 8: Supplementary Figure 8

#### Additional File 8: Supplementary Figure 8

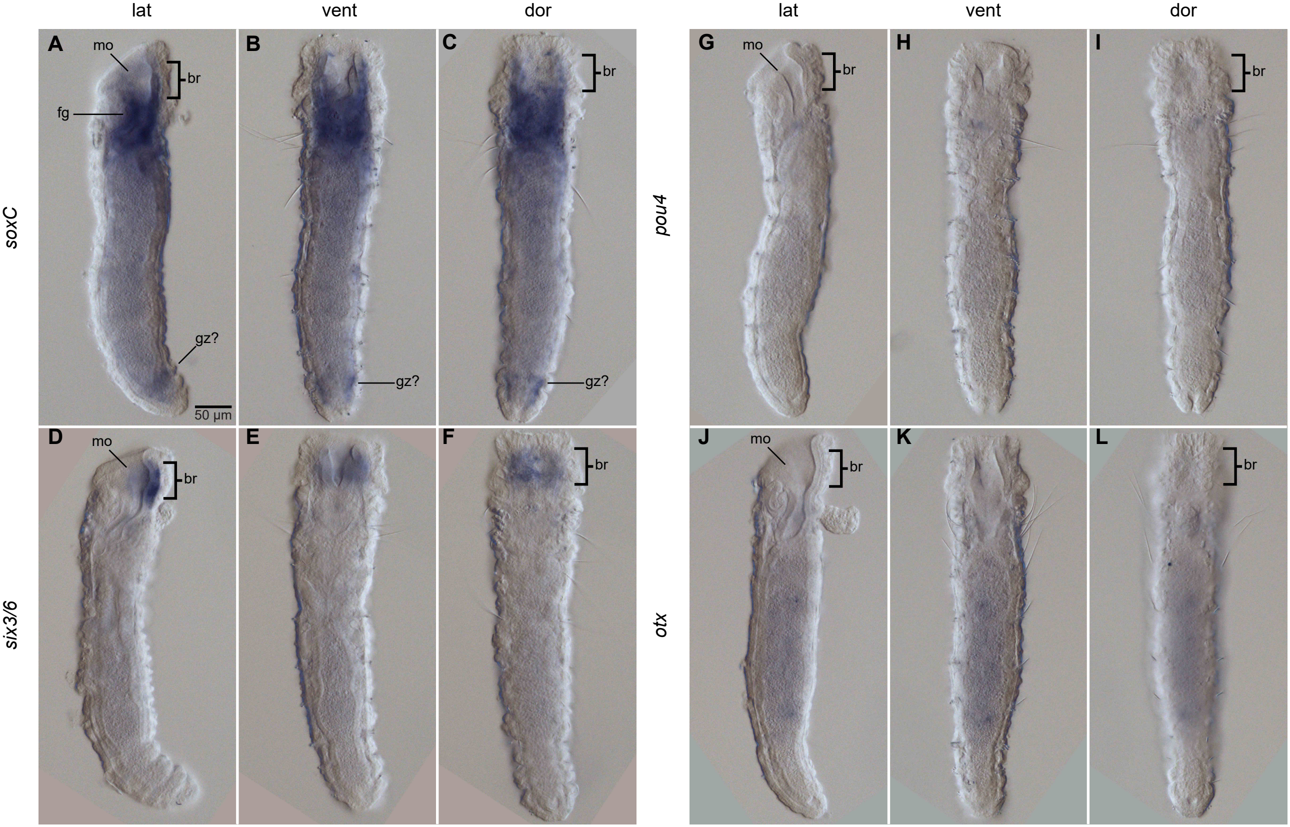

### Additional File 9: Supplementary Figure 9

#### Additional File: Supplementary Figure 9

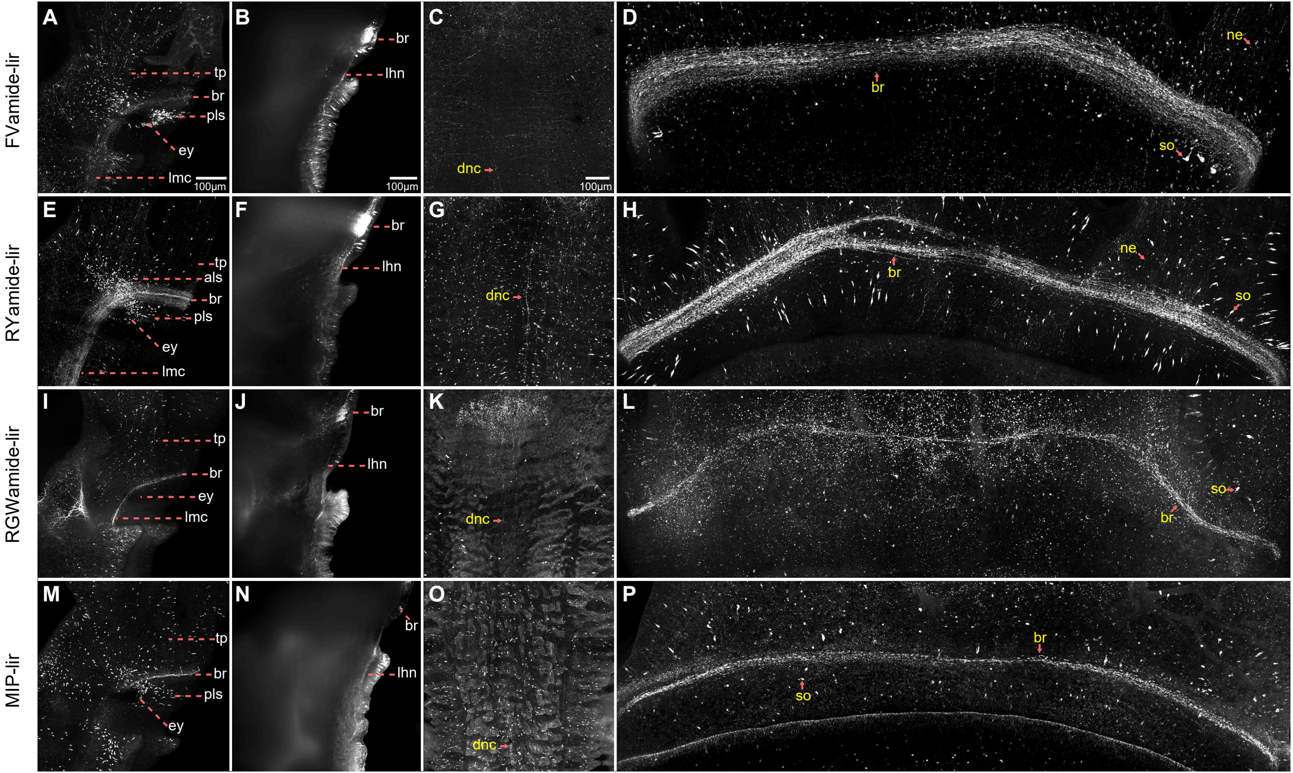
